# Direct reprogramming of human fibroblasts into insulin-producing cells by transcription factors

**DOI:** 10.1101/2021.08.05.455196

**Authors:** Marta Fontcuberta-PiSunyer, Ainhoa García-Alamán, Èlia Prades, Noèlia Téllez, Hugo Figueiredo, Rebeca Fernandez-Ruiz, Sara Cervantes, Carlos Enrich, Laura Clua, Javier Ramón-Azcón, Christophe Broca, Anne Wojtusciszyn, Anna Novials, Nuria Montserrat, Josep Vidal, Ramon Gomis, Rosa Gasa

## Abstract

Direct lineage reprogramming of one somatic cell into another bypassing an intermediate pluripotent state has emerged as an alternative to embryonic or induced pluripotent stem cell differentiation to generate clinically relevant cell types. One cell type of clinical interest is the pancreatic β cell that secretes insulin and whose loss and/or dysfunction leads to diabetes. Generation of functional β-like cells from developmentally related somatic cell types (pancreas, liver, gut) has been achieved via enforced expression of defined sets of transcription factors. However, clinical applicability of these findings is challenging because the starting cell types are not easily obtainable. Skin fibroblasts are accessible and easily manipulated cells that could be a better option, but available studies indicate that their competence to give rise to β cells through similar direct reprogramming approaches is limited. Here, using human skin fibroblasts and a protocol that ensures high and consistent expression of adenovirus-encoded reprogramming factors, we show that the transcription factor cocktail consisting of Pdx1, Ngn3, MafA, Pax4 and Nkx2-2 activates key β cell genes and down-regulates the fibroblast transcriptional program. The converted cells produce insulin and exhibit intracellular calcium responses to glucose and/or membrane depolarization. Furthermore, they secrete insulin in response to glucose *in vitro* and after transplantation *in vivo*. These findings demonstrate that transcription factor-mediated direct reprogramming of human fibroblasts is a feasible strategy to generate insulin-producing cells.

## INTRODUCTION

The discovery that adult somatic cells can be reprogrammed into induced pluripotent stem cells (iPSC) ^1, 2^ has revolutionized current biological and medical research. Human iPSC can provide valuable platforms for drug and toxicology screens, for investigating human development and for disease modeling. Furthermore, the possibility of differentiating iPSC into clinically relevant cell types for autologous cell therapy has pushed the field of regenerative medicine forward. However, current iPSC technology is still technically demanding, expensive, time consuming and produces cells of variable quality. Similarly, design of the *in vitro* differentiation protocols to generate specific cell types is not trivial. Recapitulation of in vivo developmental pathways using signaling molecules is considered the most effective approach, but defining and adjusting proper in vitro differentiation conditions is not a straightforward task, and differentiation protocols are often long and expensive, present poor reproducibility and have variable efficiencies among laboratories.

An alternative strategy to produce cell types of interest is direct lineage reprogramming, which entails the direct conversion of one differentiated cell type into another bypassing an intermediate pluripotent stage. This strategy is often based on the forced expression of cocktails of transcription factors (TF) that normally function as potent fate determinants of the cell type of interest in development ^3-5^. This approach presents several advantages over pluripotent cell derivation including its relative simplicity and speed (fewer induction steps) and, in terms of translational potential, its safety due to avoidance of the tumorigenic risk associated with pluripotency ^6^. Moreover, as with iPSC, this strategy allows autologous transplantation thus eluding the problem of allograft rejection.

Pancreatic beta (β) cells produce insulin, which controls whole body glucose homeostasis. Relative or complete deficit of functioning β cells is a hallmark of diabetes and thus β-cell replacement has emerged as a promising therapeutic approach to treat and eventually cure this disease. To date, derivation of β-cells from pluripotent (embryonic and iPS) cells has been intensively studied, with significant progress made during the past fifteen years ^7-11^. Yet, current protocols are still on the way of standardization and face important challenges including low number and functional maturation of generated β-like cells ^12, 13^. Direct lineage reprogramming strategies to create β-cells bypassing the iPSC stage have also been explored. In this line, the Melton laboratory developed the first TF-based *in situ* conversion of mouse pancreatic exocrine cells towards insulin-producing cells through expression of three TFs: Pdx1, Ngn3 and MafA (hereafter PNM)^14^. Subsequently, these factors, individually or combined, have been shown to promote β-cell-like features in related endodermal cell lineages such as pancreatic ductal cells, hepatic, intestinal or stomach cells ^15-21^.

One central aspect in direct reprogramming approaches, particularly when considering their clinical applicability, is the choice of cell source. Ideally, the starting material should be accessible, easy to handle/grow and susceptible to reprogramming. To date fibroblasts, which can be readily obtained from small skin biopsies, are commonly used as cell source for IPSC generation. Remarkably, they have also been directly reprogrammed towards somatic cell types including cardiomyocytes ^22^, chondrocytes^23^, neurons ^24^ or oligodendrocyte progenitors ^25^. In the pancreas field, human fibroblasts have been converted into expandable pancreatic progenitor populations through transient overexpression of iPSC factors in conjunction with lineage-determining soluble factors ^26^ and into insulin-producing cells following long protocols that are reminiscent of the small molecule-based directed differentiation procedures applied to stem cells ^26-28^. However, attempts to use cell-specific TFs to directly transform fibroblasts into β cells have yielded discouraging results ^14, 20, 29, 30^.

The growing potential of direct lineage reprogramming to generate specialized cell types, the medical interest in pancreatic β cells and the prevailing challenges for the generation of these cells via conventional directed differentiation from pluripotent cells, prompted us to examine the amenability of fibroblasts to be reprogrammed to β cells using cell-specific transcription factors in a more comprehensive manner. Here we show that the combination of five endocrine TF induces activation of the β-cell transcriptional program in human fibroblasts. Furthermore, we show that reprogrammed fibroblasts produce and secrete insulin *in vitro* and *in vivo*. We believe these findings demonstrate the feasibility of this approach and set the basis to explore this path for generation of insulin-producing cells for disease modeling and cellular therapy.

## MATERIALS and METHODS

### Fibroblasts

Human foreskin fibroblasts were prepared from a foreskin biopsy from a 3-year old individual after obtaining his parents’ written informed consent (HFF1). Adult human dermal fibroblasts were prepared from arm biopsies of two T1D individuals after obtaining their written informed consent (HDF-DT1-1, HDF-DT1-2). In brief, skin samples were collected in sterile saline solution, divided into small pieces, and allowed to attach to cell culture dishes before adding Iscove’s modified Dulbecco’s medium (Invitrogen, Carlsbad, CA, USA) supplemented with 10% human serum (Sigma, St. Louis, MO, USA) and penicillin/streptomycin (0.5X) (Invitrogen). After 10 days of culture at 37°C, 5% CO2, fibroblast outgrowths were dissociated and split 1:4 using a recombinant trypsin-like enzyme (TrypLE Select, Invitrogen). Additional fibroblast preparations were commercially available: HFF2 (SCRC1041™, ATCC, Manassas, VA, USA) and HDF-1 (PCS-201-012, ATCC) and HDF-2 (Cell Applications, San Diego, CA, USA).

### Human islets

Human islets were prepared by collagenase digestion followed by density gradient purification at the Laboratory of Cell Therapy for Diabetes (Hospital Saint-Eloi, Montpellier, France), as previously described ^31^. After reception in Barcelona, human islets were maintained in culture for 1-3 days in RPMI-1640 with 5.5mM glucose, 10% fetal bovine serum (FBS) and antibiotics, before performing the experiments. Experiments were performed in agreement with the local ethic committee (CHU, Montpellier) and the institutional ethical committee of the French Agence de la Biomédecine (DC Nos. 2014-2473 and 2016-2716). Informed consent was obtained for all donors.

### Recombinant adenoviruses

The adenoviral expression vector pAd/CMV/V5-DEST carrying mouse Pdx1, Ngn3, MafA and 2A-Cherry under the CMV promoter was kindly provided by Dr. Q. Zhou, Harvard University^29^. The recombinant adenovirus (termed Ad-PNM) was generated after Pac1 digestion and transfection into HEK293 cells. The recombinant adenovirus encoding Pdx1 was kindly provided by the Beta Cell Biology Consortium. The recombinant adenovirus encoding MafA was purchased from Vector Biolabs (Chicago, IL, USA). All other recombinant adenoviruses encoding single transcription factors (Ngn3, NeuroD1, Nkx2.2, Nkx6.1 and Pax4) were described previously^18, 32^.

### Reprogramming protocol

Fibroblasts were grown in DMEM media supplemented with 10% (v/v) fetal bovine serum (FBS), 100 U/ml penicillin, 100 µg/ml streptomycin and 1% Glutamax. They were plated onto 96-well plates (9 500 cells per well) for MTT and BrdU assays, onto 12-well plates (1.25×10^5^ cells/well) for gene expression, insulin secretion, immunofluorescence and caspase assays and onto 10 cm plates (1.5×10^6^ cells/plate) for transplantation experiments. Infection with Ad-PNM was performed when fibroblasts reached 80% confluence, normally 1-2 days after seeding. In brief, for 12-well plates, fibroblasts were incubated with 0.6ml complete culture medium containing Ad-PNM at a dose of 15 moi (multiplicity of infection) for 16-20 h. After virus removal, media was changed to RPMI-1640 medium containing 6% FBS and antibiotics. Two and five days after incubation with Ad-PNM, fibroblasts were infected for 6-8 h with Ad-Pax4 (50 moi) and Ad-Nkx2.2 (50 moi) respectively. The transfection reagent Superfect (Qiagen, Venlo, Netherlands) was always added to the virus-containing media in order to achieve efficient adenoviral infection and transgene expression in fibroblasts. Amounts were scaled down or up according to the well size. To prepare spheroids, one day after Ad-Nkx2.2 infection, cells were trypsinized and transferred (1 200-1 800 cells/ well) to 96-well Nunclon Sphera plates (Thermo Scientific) for gene expression, immunofluorescence and insulin secretion assays. Cell spheroids (1 000 cells/cluster) for transplantation were generated in AggreWell plates (StemCell Technologies, Saint Égrève, France)

### RNA isolation and reverse transcriptase polymerase chain reaction (RT-PCR)

Total RNA from cultured cells was isolated using NucleoSpin®RNA (Macherey-Nagel), Düren, Germany) following the manufacturer’s manual. Total RNA from enucleated eyes was extracted using Trizol reagent (Sigma) and then cleaned and DNAse-treated using RNeasy mini columns (Qiagen) prior to cDNA synthesis. First-strand cDNA was prepared using Superscript III Reverse Transcriptase (Invitrogen) and random hexamers in a total volume of 20 ul and 1/40 to 1/200 of the resulting cDNA was used as a template for real time PCR reactions. Real time PCR was performed on an ABI Prism 7900 detection system using Gotaq master mix (Promega, Madison, WI, USA). Expression relative to the housekeeping gene *TBP* was calculated using the delta(d)Ct method and expressed as 2^(-dCT) unless otherwise indicated. Primer sequences are provided in Table S3.

### BrdU and MTT assays

For quantification of cell proliferation, cells in 96-well plates were cultured overnight with medium containing 5-bromo-2’-deoxuridine (BrdU). BrdU incorporation was determined colorimetrically with the Cell proliferation ELISA kit (Roche, Basilea, Switzerland) following the manufacturer’s instructions. For assessing cell viability, cells grown in 96-well plates were incubated with medium containing 0.75 mg/ml of 3-(4,5-dimethythiazol-2-yl)-2,5-diphenyltetrazolium bromide (MTT) for 3 hours at 37ºC. The resulting formazan crystals were solubilized in Isopropanol/0.04 N HCl solution and optical density ^1^ was read at 575 nm and 650 nm using a Synergy HT reader (BIO-TEK Instruments, Winooski, VT, USA). The OD (575-650) was expressed relative to control fibroblasts, which were given the value of 100%.

### Calcium imaging

To study glucose-dependent calcium influx, cells were washed with Hanks’ Balanced Salt Solution (HBSS, Sigma) and incubated with Fluo4-AM (Life Technologies) in fresh 2.5 mM glucose Krebs-Ringer bicarbonate buffer for 1 hour at 37ºC in the dark. Intracellular calcium fluorescence was recorded from fluo-4-loaded cells using a Leica TCS SPE confocal microscope with an incubation chamber set at 37C, and a 40x oil immersion objective. Fluorescent images and average fluorescence intensity were acquired at 600Hz every 1.8 seconds, using a 488nm excitation laser, an emission set at 520 with a bandwidth of 10nm. Image registry consisted of: 5 min in 2 mM glucose-Krb buffer, 10 min in 22 mM glucose-Krb buffer and 5 min in 30 mM KCl-Krb buffer. Average fluorescence intensity images of each individual cell were analyzed with LAS AF Lite and Fuji programs.

### In vitro glucose-induced insulin secretion and content

Cells in 2D or 3D aggregates were washed with phosphate-buffered saline (PBS) and preincubated for 30-45 min at 37°C in Hepes-buffered Krebs-Ringer buffer with 2mM glucose. Preincubation solution was then removed and cells incubated with the same buffer containing different glucose concentrations glucose for additional 90 min. Supernatants were collected to measure secreted insulin or C-peptide. Cells were lysed in detergent-containing buffer (Tris-HCl pH8 50mM, NaCl 150mM, SDS 0.1% (w/v), Igepal CA-630 1%, Na deoxycholate 0.5%) to determine C-peptide content. Human insulin was measured using a human Insulin ELISA kit (Crystal Chem, Zaandam, Netherlands) and C-peptide using a human C-peptide ELISA kit (Mercodia, Uppsala, Sweden).

### Electron microscopy

Cell spheroids were collected, washed with PBS and fixed with 4% paraformaldehyde/ 0.5% glutaraldehyde (Sigma) mixture in 0.1M phosphate buffer (PB) pH7.4 for 30min at 4°C and gentle agitation. Cells were then transferred to fresh fixation solution and maintained at 4°C until secondarily fixed with 1% uranyl acetate and 1% osmium tetroxide. Cells were then dehydrated, embedded in Spurr’s Resin and sectioned using Leica ultramicrotome (Leica Microsystems). Conventional transmission electron microscopy (TEM) images were acquired from thin sections using a JEOL-1010 electron microscopy equipped with an SC1000 ORIUS-CCD digital camara (Gatan).

### Transplant procedures

Adult (12-20 week old) immunocompromised NSG-SCID mice (005557, Jackson Laboratories) received implants in the anterior chamber of the eye (ACE), subcutaneously or between the omental sheets as previously described^33, 34^. A total of 150-200 human islets, 300 iβ-like cell or HFF1 spheroids (1 200-1 800 cells/cluster) were transplanted per eye and approximately 4000-iβ-like cell spheroids (750-1 000 cells/cluster) were transplanted into either of the other two implant sites. For subcutaneous and omental transplants, collagen scaffolds were used. Briefly, the cell-laden collagen hydrogel was prepared by mixing 150 μl of a 4 mg/ml rat tail type I collagen (Corning, NY, USA) solution, prepared following the manufacturer’s instructions, with iβ-like cell clusters (pelleted by gentle centrifugation) and poured onto a cylindrical 8 mm diameter x 1 mm thick PDMS mold (Dow Corning Sylgard 184 Silicone Elastomer). The hydrogel was polymerized for 20 min at 37 ºC, detached from the mould and maintained in tissue culture dishes with warm RPMI-1640 medium until transplant (usually 2-3 hours). Constructs were introduced in the abdominal subcutaneous space through a small (1–2 mm) incision. Alternatively, constructs were placed on the omentum close to the duodenal-stomack junction and fixed using Histoacryl translucent (Braun, Kronberg im Taurus, Germany). The local ethical committee for animal experimentation of the University of Barcelona approved all animal experiments and procedures.

### Graft viability and function analysis

To assess vascularization and cell viability in ACE implants, iβ-like clusters were labeled with the long-term tracer for viable cells CFDA (Invitrogen) before transplantation. At day 10 post-transplantation, mice received an intravenous injection of RITC-dextran and in vivo imaging was used to assess functional vascularization and cell viability ^34^. For determination of glucose-induced insulin secretion, mice were fasted for 5-6 h and then injected intraperitoneally with glucose (3g/Kg). Aqueous humor from mice with ACE implants was obtained 30 minutes after the glucose challenge and kept frozen until human insulin determination. Tail blood before and after (20 minutes) the glucose challenge was obtained from mice with subcutaneous and omental implants. Human insulin in aqueous humor and in plasma was determined using an ultrasensitive Human Insulin ELISA (Chrystal Chem).

### Immunofluorescence

iβ-like cells grown in 2D were fixed with 4% (v/v) paraformaldehyde (PFA) during 15 min and incubated with blocking solution (0.25% (v/v) Triton, 6% (v/v) donkey serum, 5% (w/v) BSA in PBS for 1 h at room temperature. Slides were then incubated with primary antibodies diluted in PBS-triton 0.1% (v/v) containing 1% donkey serum overnight at 4 o C. iβ-like cells grown in spheroids were fixed with 4% (v/v) paraformaldehyde (PFA) for 15 min at 4ºC, permeabilized with 0.5% (v/v) Triton in PBS for 20 min and blocked with 0.5% (v/v) Triton/ FBS 10% (v/v) in PBS during 1 h at room temperature. Slides were then incubated with primary antibodies diluted in blocking solution overnight at 4°C. Eyes were obtained by enucleation, fixed overnight in 2% (v/v) PFA, dehydrated with ethanol gradient, cleared with xylene and paraffin-embedded. 3-µm thick eye sections were used for standard immunofluorescence staining protocol. Images were acquired in a Leica DMR HC epifluorescence microscope (Leica Microsystems). Primary antibodies used were: Insulin (DAKO, 1:400), C-peptide (Abcam, 1: 200 or Hybridoma Bank; 1:40), vimentin (Sigma, 1:200), HLA (Abcam, 1:100), Nkx-2.2 (Hybridoma Bank, 1:400), Pdx1 (Abcam 1:500) and MafA (Novus Biologicals 1:75). The antigen-primary antibody immune complex was visualized with secondary antibodies conjugated to Alexa Fluor 488 (Jackson Immunoresearch, 1:250), Alexa fluor 555 (Molecular Probes, 1:400) or Alexa fluor 647 (Jackson Immunoreserach, 1:250). Cell nuclei were counterstained with Hoescht (SIGMA, 1:500). Representative images were taken using a confocal microscope (LEICA TCS SPE).

### Statistical analysis

Data are presented as mean ± standard error of the mean (SEM) from at least three independent reprogramming experiments, with one to four biological replicates per experiment. Significant differences between the means were analyzed by the two-tailed unpaired Student’s *t*-test, one sample t-test or one-way ANOVA followed by Tukey’s multiple comparison tests as indicated in the figure legends. Statistical analysis was performed with GraphPad Prism 8.00 and Microsoft Office Excel 2007 and differences were considered significant at p<0.05. No methods were used to determine whether the data met assumptions of the statistical approach (e.g. test for normal distribution).

## RESULTS

### Introduction of the transcription factors Pdx1, Ngn3 and MafA (PNM) leads to activation of β-cell genes and insulin production by human fibroblasts

We first investigated whether human fibroblasts could be directly converted towards the β-cell lineage through exogenous expression of the PNM factors Pdx1, Neurog3 and MafA. As delivery method we used a polycistronic adenoviral vector encoding the three transcription factors and a Cherry reporter protein (Ad-PNM hereafter) that had been previously used in the reprogramming of pancreatic acinar cells towards islet cells ^35^. We chose adenoviruses for transgene delivery because they allow high expression of transgenic proteins and, most importantly, don’t integrate in the host genome. However, we couldn’t detect Cherry immunofluorescence after viral treatment of human fibroblasts (Fig. S1), which was in agreement with other reports showing poor adenoviral transduction efficiency in fibroblastic cells ^36, 37^. In an attempt to boost adenoviral transduction, we added an activated dendrimer normally used in transfections to the virus-containing media and, in stark contrast to Ad-PNM alone, we observed abundant (>80%) fibroblasts that were Cherry^+^ three days after exposure to the virus+dendrimer (Fig.S1 and Fig.1A). Interestingly, Cherry^+^ cells were still readily visible 7 days post-viral treatment (Fig. 1A). Likewise, transcripts encoding the PNM factors were detected at 3 and 7 days after viral treatment (Fig.1B). Remarkably, three days after addition of Ad-PNM we detected low levels of *INS* transcripts that were dramatically increased by day 7 (Fig.1C). We examined the influence of distinct culture media on *INS* mRNA levels and found that cells cultured in RPMI-1640 supplemented with 6% FBS exhibited the highest degree of *INS* gene activation relative to cells cultured in DMEM or CMRL-1066, reaching values that were 900-fold lower than human islets but similar to those attained in directed differentiation protocols from iPSC ^38^ (Fig. 1D). The PNM factors also induced moderate levels of several endocrine transcription factor genes including *NEUROD1, INSM1, PAX4, NKX2-2* and *ARX*, although they did not induce the key β-cell TF *NKX6-1* (Fig.1E). Unexpectedly, the PNM factors also induced the islet hormone genes *GLUCAGON (GCG)* and *SOMATOSTATIN (SST)*, albeit to levels less than those observed for *INS (*as indicated by lower expression relative to the housekeeping gene *TATA BINDING PROTEIN*, (*TBP*) (Fig. 1F). Conversely, PNM down-regulated the expression of genes associated with fibroblast signature, including several factors involved in maintenance of the fibroblastic transcriptional program such as *TWIST2, PRRX1* and *LHX9* ^39^ (Fig. 1G). Together, these experiments show that the PNM factors induce a transcriptional switch from a fibroblastic towards an endocrine program in human fibroblasts.

**Figure 1.**
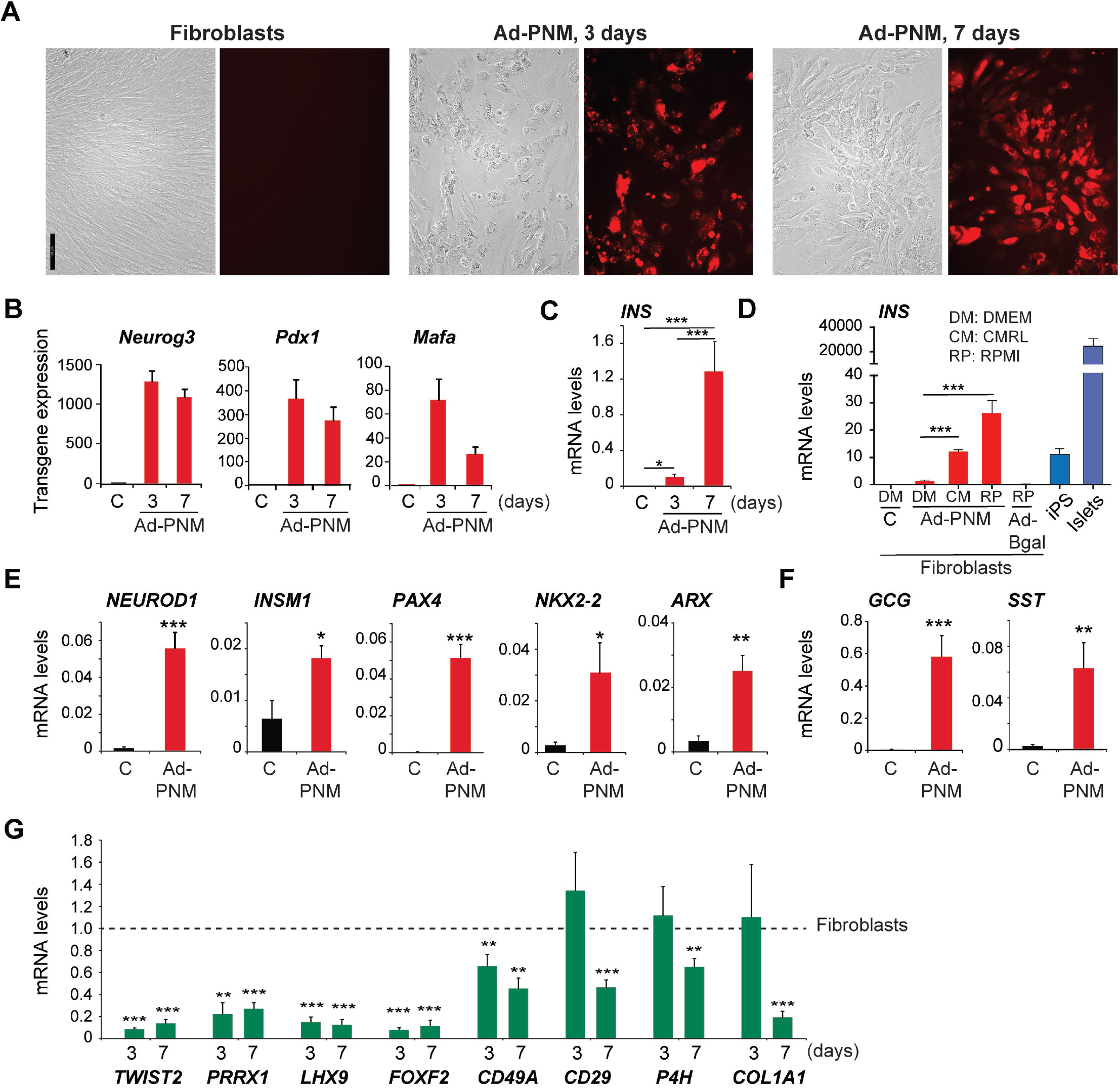
Introduction of the PNM factors in human fibroblasts. Human foreskin fibroblasts (HFF1) were transduced with a polycistronic recombinant adenovirus encoding the transcription factors Pdx1, Neurog3, MafA and the reporter protein Cherry (Ad-PNM). Untreated parental fibroblasts were used as controls (indicated as C in graphs). **(A)** Bright field images and Cherry immunofluorescence of control fibroblasts and fibroblasts infected with Ad-PNM at day 3 and 7 post-infection. Scale bar is 100 µm. **(B)** qRT-PCR of adenovirus-encoded transgenes at day 3 (n=11) and 7 (n=6) after infection with Ad-PNM. **(C)** qRT-PCR of human *INS* at day 3 (n=7) and 7 (n=13) after infection with Ad-PNM. (**D)** In red, qRT-PCR of human *INS* in fibroblasts maintained in the indicated culture media during 7 days after infection with the indicated adenoviruses, (n=3-10). In blue, *INS* mRNA levels in iPSC-derived β-cells and in isolated human islets (n=10). **(E-F)** qRT-PCR of indicated genes encoding islet/β-cell transcription factors and islet hormones at day 7 post-PNM (n=8-15). **(G)** qRT-PCR for the indicated fibroblast markers at days 3 (n=5-11) and 7 post-PNM (n=5-6). In **B-F**, expression levels are expressed relative to *TBP*. In ***G***, expression is relative to control fibroblasts, given the value of 1 (dotted line). Data are presented as the mean ± SEM for the number of samples indicated in parenthesis. *,P<0.05; **,P<0.01; ***,P<0.001, relative to control (untreated) fibroblasts or between indicated conditions using unpaired t-test (**B-E**) or one sample t-test (**G**).

### Addition of Pax4 and Nkx2-2 to the PNM factors improves reprogramming of human fibroblasts towards the β-cell lineage

Due to the suboptimal conversion of fibroblasts into β-like cells using the PNM factors (i.e. as reflected by the lack of induction of *NKX6-1* mRNA and the expression of other endocrine hormones), we tested the effects of adding other transcription factors to the reprogramming cocktail, namely Nkx6-1, Pax4, NeuroD1 and Nkx2-2. Among all tested combinations, we found that expression of PNM in a single polycistronic adenovirus followed by sequential expression of Pax4 and Nkx2-2 (hereafter this combination is named 5TF) resulted, ten days after initial PNM infection, in the highest degree of *INS* gene activation accompanied by the blockade of *GCG* gene induction (Fig. 2A&B and Fig.S2). Moreover, 5TF reprogramming resulted in activation of *NKX6*.*1* and of the pan-endocrine factor *PAX6 genes (*Fig. 2B). At day 10, reprogrammed cells displayed an epithelial morphology (Fig. 2C) and hadn’t grown as much as untreated fibroblasts (day 10; 5TF: 44×10^3^ ± 3×10^3^ cells/well; control: 238×10^3^ ± 18×10^3^ cells/well, n=18). In fact, reprogrammed cells showed diminished proliferation as early as one day after PNM introduction, while remaining viable, as indicated by their retained ability to reduce MTT, similar to that of fibroblasts (Fig. 2D&E). At day 10, reprogrammed cells exhibited lower MTT reduction activity, likely due to their decreased cell numbers (Fig. 2E).

**Figure 2.**
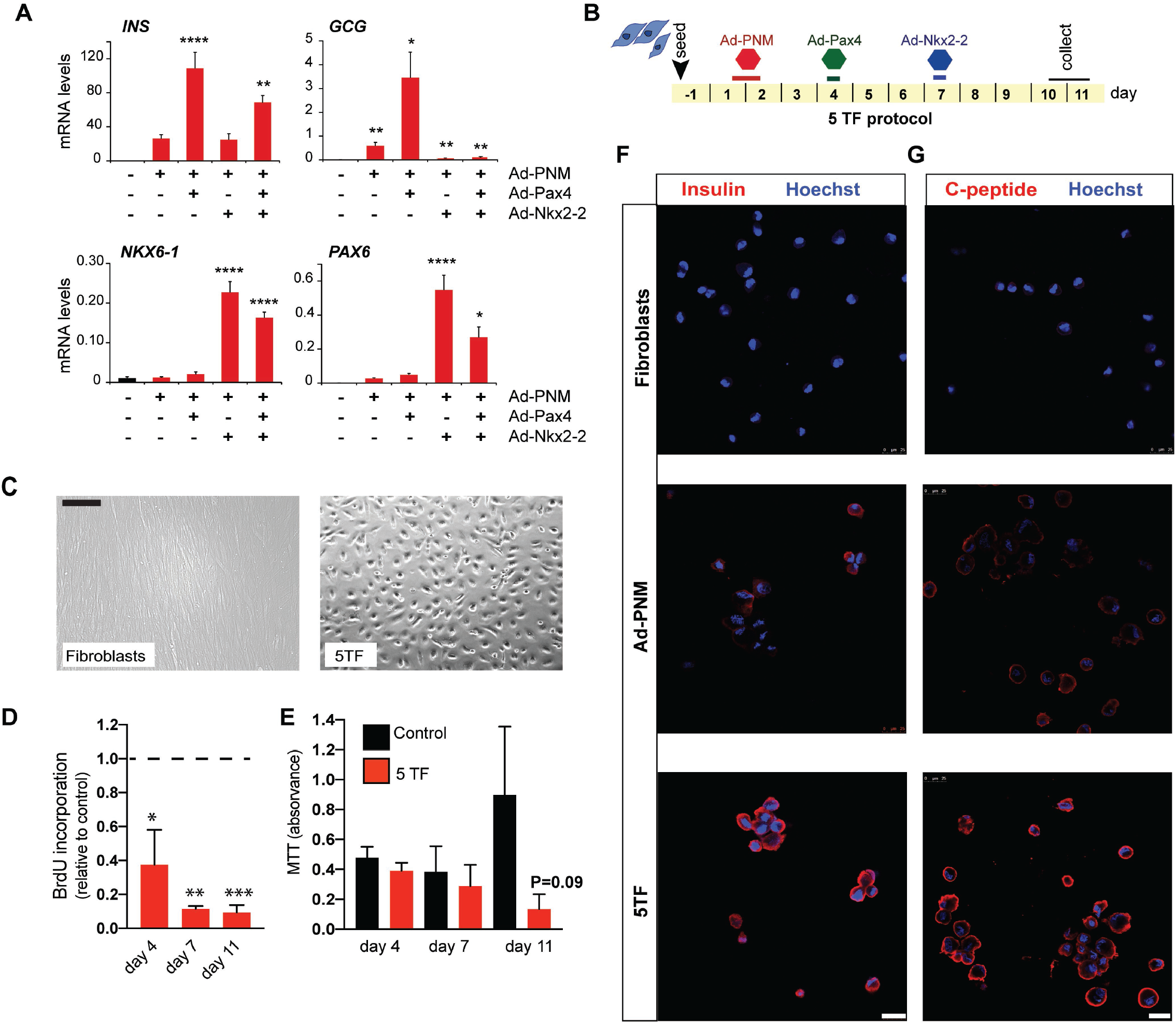
Sequential introduction of the PNM factors, Pax4 and Nkx2-2 (5 TF protocol) in human fibroblasts. **(A)** qRT-PCR for the indicated genes in HFF1 fibroblasts infected with different combinations of the indicated adenoviruses. Pax4 was added three days after PNM. Nkx2-2 was added three days (PNM+Nkx2-2) or six days (PNM+Pax4+Nkx2-2) after PNM. Expression levels are calculated relative to *TBP* (n=4-12). **(B)** Scheme of the final 5TF reprogramming protocol showing the sequence of addition of adenoviruses encoding the indicated transcription factor/s. Duration of incubation with each adenovirus is represented with a line. Cells were studied at days 9-10 after initial addition of Ad-PNM. **(C)** Representative bright field image of parental fibroblasts and 5TF reprogrammed fibroblasts at day 10. **(D)** Cell proliferation measured by BrdU incorporation (n=3) and **(E)** Cell viability measured by MTT assay (n=3) in untreated control (in black) and 5TF-reprogrammed fibroblasts (in red) at the indicated days. Day 4 values are before Pax4 introduction. **(F-G)** Representative immunofluorescence images showing Insulin and C-peptide staining (in red) in untreated fibroblasts and in fibroblasts infected with Ad-PNM alone or with 5TF. Nuclei were stained with Hoechst (in blue). Scale bar is 25 μm. Data are presented as the mean ± SEM for the number of experiments indicated in parenthesis *,P<0.05; **,P<0.01; ***,P<0.001, P<0.0001 relative to PNM (in **A**) or to control fibroblasts (**D**,**E**) using one-way ANOVA (**A**), one-sample t-test (**D**) or unpaired t-test (**E**).

Consistent with the gene expression data, insulin protein was detected by immunofluorescence staining in both PNM and 5TF cells, being staining more robust in 5TF cells than in PNM cells (Fig. 2F). By contrast, neither glucagon nor somatostatin proteins were detected in either PNM or 5TF cells. We quantified the immunofluorescence images and found that 67.9±6.2% of cells in the culture were INS+ at day 10. The presence of endogenous insulin production was further confirmed by immunofluorescence staining using an antibody against C-peptide (Fig.2G) and by determining C-peptide content with an ELISA (control: 0.34 ± 0.14 pg/10^5^ cells, n=9; 5TF: 16.06 ± 2.55 pg/10^5^cells, n=16). To further establish the extent of reprogramming achieved with 5TF as compared to PNM, we studied expression of additional endocrine genes at day 10. We found that transcripts of β-cell differentiation TFs (i.e. *NEUROD1, INSM1, HNF1B, MAFB*), as well as some of the reprogramming TFs (*PDX1, NEUROG3, NKX2-2*) were higher in 5TF as compared to PNM (Fig. 3A). On the other hand, several genes associated with a mature β-cell phenotype were more upregulated by 5TF than PNM (*PCSK1, ABCC8, KCNJ11, GLP1R, GIPR, NCAM1, CXCR4*), whilst others (*CHGB, PTPRN, CX36*) were induced to similar extents by both TF combinations (Fig. 3B). In line with loss of *GCG* activation, the pro-convertase gene *PCSK2*, which is expressed at higher levels in α than β cells ^40^, was reduced by 5TF as compared to PNM (Fig.3B), supporting that sequential introduction of Pax4 and Nkx2-2 after PNM endorses the β-cell differentiation program in human fibroblasts. It should be noted that expression levels of the induced genes were lower than in human islets, with differences ranging from 2-fold for *NCAM1* to 20,000-fold for *CHGB* (Fig. 3A,B, Table S1). Gene activation was sustained for at least eleven additional days under the same culture conditions despite declined expression of the reprogramming factor transgenes. Moreover, expression of some genes increased with time in culture (i.e. *NKX6*.*1, INSM1, ABCC8, CHGB*) (Fig.S3).

**Figure 3.**
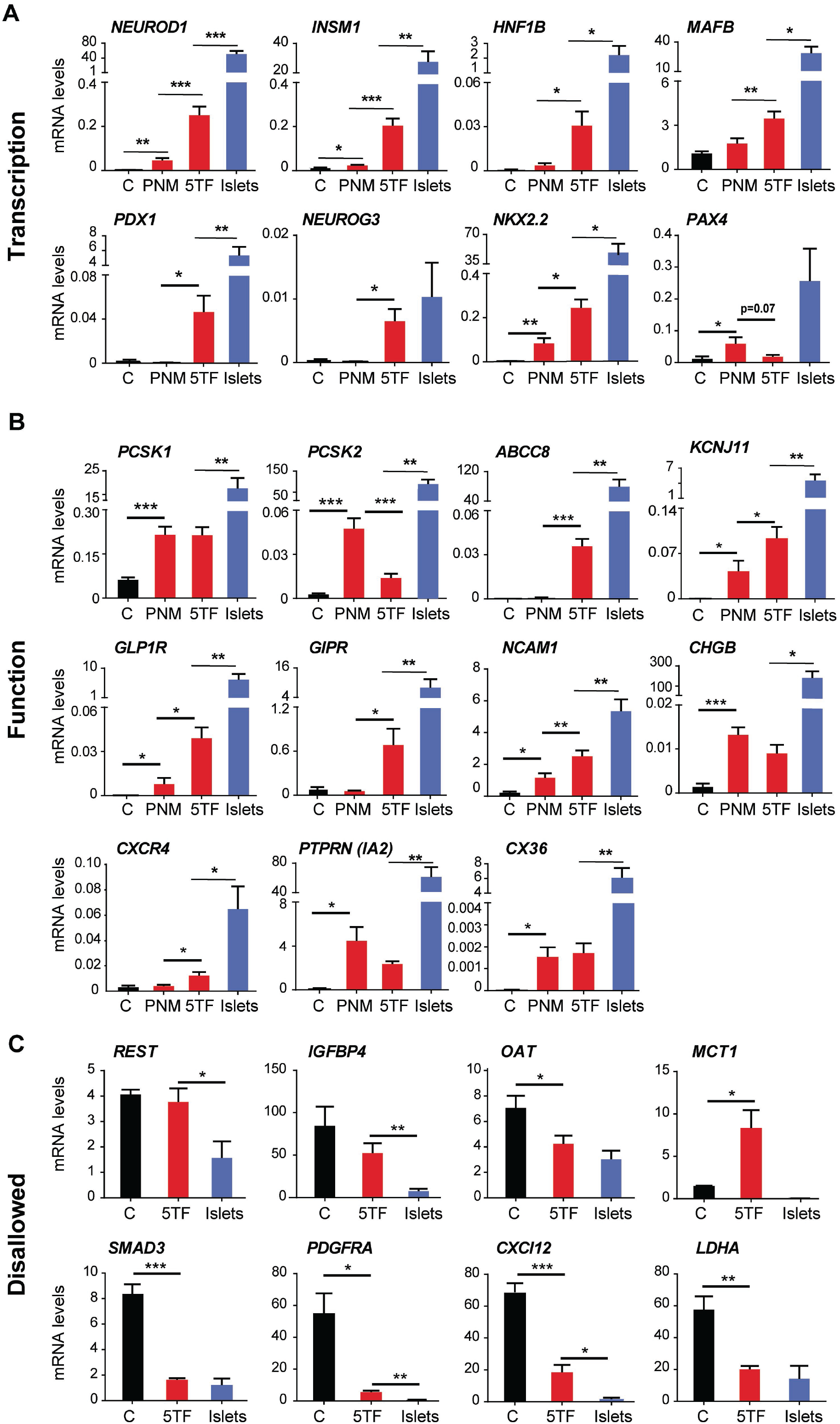
5FT protocol promotes β-cell specific gene activation and repression events in human fibroblasts. **(A**,**B)** qRT-PCR of islet/β-cell transcription factor **(A)** and β-cell function **(B)** genes in untreated control fibroblasts (n=5-19), in fibroblasts infected with Ad-PNM alone (n=3-20) or with 5TF (n=5-18) and in human islets (n=6-9). **(C)** qRT-PCR of β-cell disallowed genes in untreated control fibroblasts (n=5), in fibroblasts infected with 5TF (n=5-8) and in human islets (n=5). Expression levels were calculated relative to *TBP*. Values represent the mean ± SEM for 5-14 for the number of n indicated in parenthesis *, P< 0.05, **, P< 0.01, ***, P< 0.001 between indicated bars using unpaired t-test.

Lastly, we looked into genes that are selectively repressed (disallowed) in β cells ^41^. In response to 5TF, several β-cell disallowed genes were significantly downregulated reaching expression levels comparable to human islets (*OAT, SMAD3, PDGFRA, CXCL12, LDHA*), while others remained unchanged (*REST*, I*GFBP4*). Among the studied genes only *MCT1* was induced by 5FT relative to parental HFF1 cells (Fig.3C).

Together, these results reveal that 5TF induces a transcriptional switch from a fibroblastic to a β-like cell identity that entails both selective gene activation and repression events. Hereafter, we will refer to cells generated following the 5TF protocol as induced β-like (iβ-like) cells.

### In vitro assessment of human fibroblast-derived iβ-like cells

Glucose-induced insulin secretion (GSIS) by β cells is mediated by glucose metabolism, closure of ATP-dependent potassium channels, membrane depolarization and opening of voltage-dependent calcium channels, resulting in an increase in cytosolic Ca^2+^ that triggers insulin exocytosis. We investigated whether iβ-like cells exhibited β-cell functional features *in vitro* and begun by determining their ability to increase intracellular Ca^2+^ in response to glucose and membrane depolarization elicited by high potassium. We found that 65% of the cells exhibited a response to glucose, high potassium, or both whilst 35% of cells were unresponsive to either stimulus (Fig. 4A and video S1). Parental HFF1 cells not engineered for 5TF expression were universally unresponsive to these stimuli (Fig. 4B and video S2). Among responsive cells, approximately half responded to both glucose and high potassium and half responded only to potassium (Fig. 4A). We observed heterogeneity in the amplitude and kinetics of responses among individual cells (Fig. 4C). Next, we performed static incubation assays to study glucose-induced insulin secretion (GSIS) and found that iβ-like cells released similar amounts of human insulin at low (2mM) and high (20mM) glucose concentrations (Fig.4D). Thus, at this stage, iβ-like cells lack a key signature feature of normal β-cells, the ability to increase insulin secretion in response to changes in extracellular glucose.

**Figure 4.**
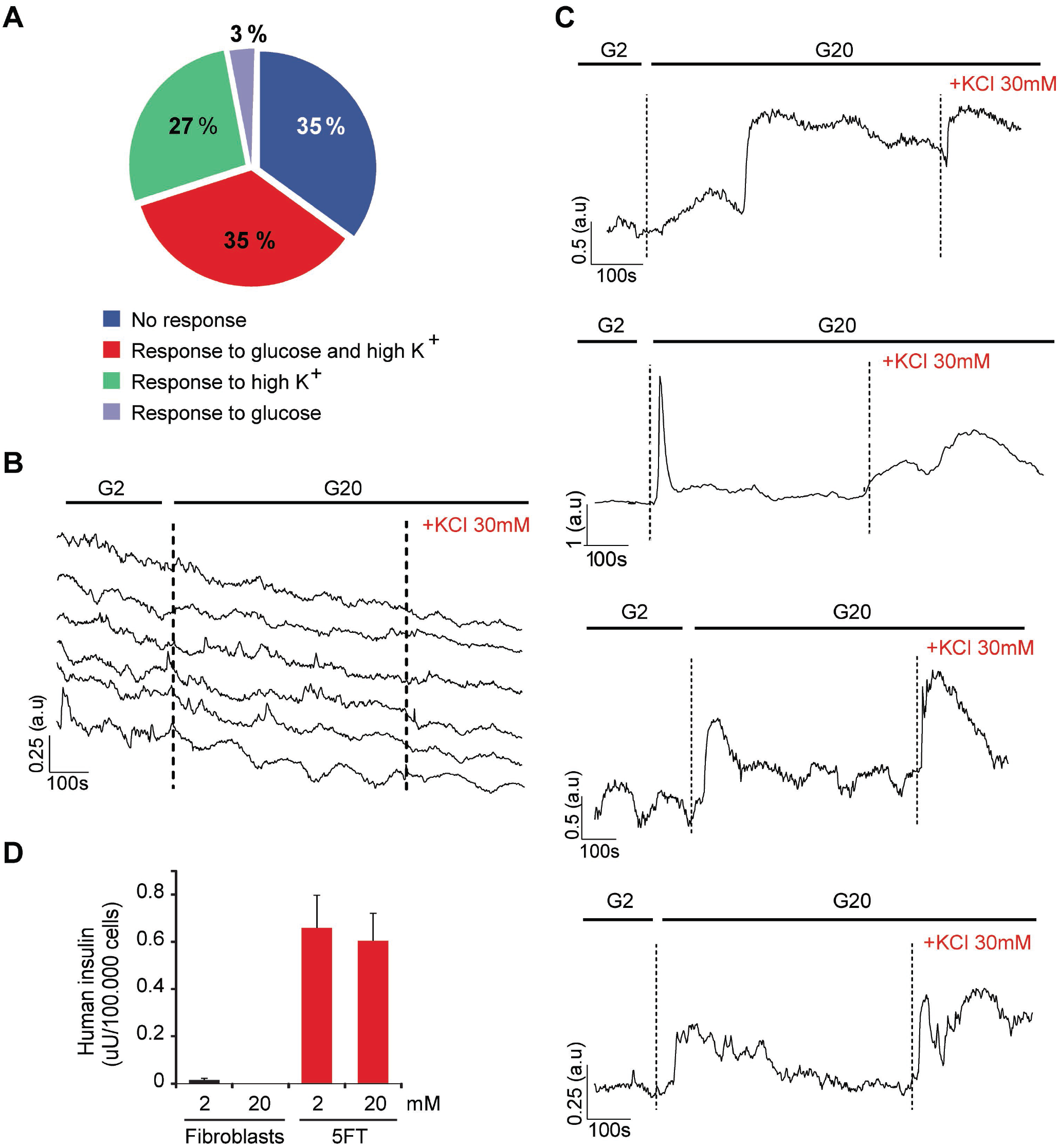
Functional characterization of iβ-like cells generated with the 5TF protocol. iβ-like cells were loaded with the calcium indicator Fluo-4-AM at day 10 of the 5TF protocol. Single-cell imaging to detect cytosolic calcium was performed in the following sequence: low glucose (2mM, G2), high glucose (20mM, G20) and membrane depolarization with KCl 30mM. **(A)** Quantification of the frequency of cells (n=200, from 6 independent reprogramming experiments) that responded to glucose, membrane depolarization elicited by high potassium or both. Representative measurements of dynamic Fluo-4 fluorescence for **(B)** six single fibroblasts and **(C)** four single iβ-like cells. **(D)** In vitro insulin secretion by iβ-like cells. ELISA determination of secreted human insulin by control fibroblasts and iβ-like cells under non-stimulatory conditions (glucose 2mM) and stimulatory conditions (glucose 20 mM). Data are mean ± SEM for 6 independent reprogramming experiments.

The differentiation and functionality of many cell types vary dramatically between three-dimensional (3D) and two-dimensional (2D) monolayer cultures, the former being closer to the natural 3D microenvironment of cells in a living organism. Considering this, we reasoned that moving from the planar cell culture to a 3D system grown in suspension might improve the functionality of iβ-like cells. To test this, we generated spheroids of 1 200-1 800 iβ-like cells (average diameter of spheroids were 128±27 µm) one day after introduction of Nkx2-2 and maintained them in culture for three additional days (Fig.5A). Insulin protein was readily detected by immunostaining (Fig. 5B). Further, iβ-like cell spheroids presented higher transcript levels for several β-cell maturation markers including *INS*, the prohormone convertase *PCSK1*, and the ATP-dependent potassium channel subunits *KCNJ11* and *ABCC8*, as compared to iβ-like cells maintained in 2D cultures (Fig.5C). Consequently, the differences in expression levels with human islets were reduced (Table S2). Consistent with improved maturation; iβ-like cell spheroids exhibited a moderate but significant insulin secretory response to glucose (fold 20mM/2mM: 2.02±0.18) as compared to 2D cultures (fold 20mM/2mM: 1.08-±0.15) in static GSIS assays (Fig. 5D,E). Similar results were obtained using a different preparation of human foreskin fibroblasts (Fig. S4). Therefore, three-dimensional culture during the last stage of the 5TF reprogramming protocol (note that total length of the protocol remained unchanged) improved iβ-like cell function

**Figure 5.**
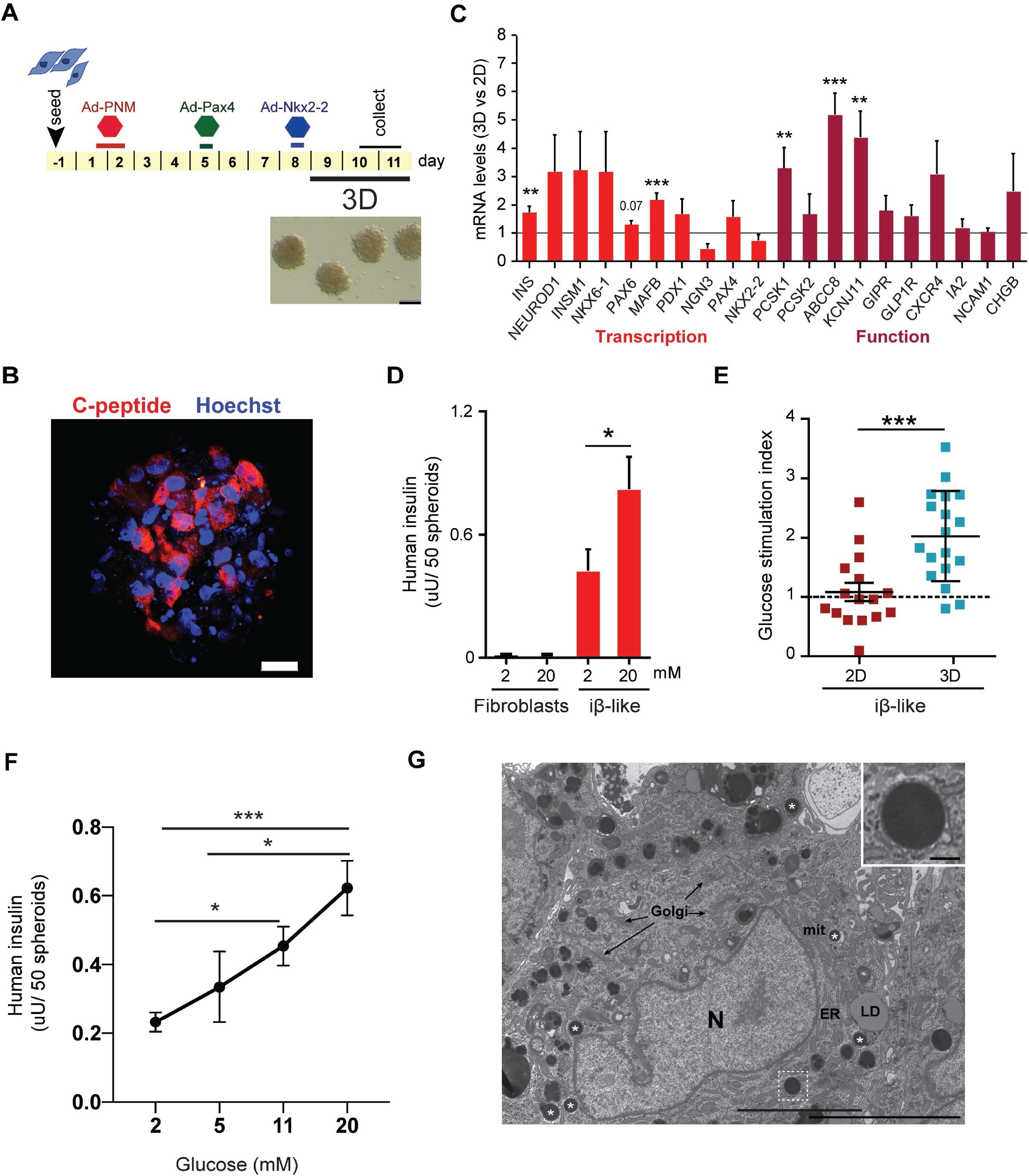
In vitro characterization of iβ-like cell spheroids. **(A)** Schematic representation of the modified 5TF protocol: cells were moved from 2D to 3D culture during the last three days (days 7-10) of the protocol. Bright field image of generated iβ-like cell spheroids. Scale bar is 100 μm. **(B)** Representative immunofluorescence image showing C-peptide staining in red and nuclei in blue (marked with Hoechst) of an iβ-like cell spheroid at the end of the reprogramming protocol. Scale bar is 100 µm. **(C)** qRT-PCR of the indicated genes in iβ-like cell spheroids (n=9-16). Transcript levels are expressed as fold relative to levels in iβ-like cells maintained in 2D culture throughout the 10-day protocol (given the value of 1, dotted line). **(D)** In vitro glucose-induced insulin secretion by iβ-like cell spheroids (n=14, from 8 reprogramming experiments). Secretion by control spheroids composed of parental fibroblasts (n= 5) is also shown. **(E)** Glucose secretory Index (fold-response 20mM vs. 2mM glucose) of iβ-like cells maintained in 2D or in 3D (spheroids) cultures (n=16-18, from 8-10 reprogramming experiments). **(F)** Glucose dose curve of insulin secretion by iβ-like cell spheroids (n=4-12, 5 reprogramming experiments). **(G)** Conventional transmission electron microscopy showing a representative image of iβ-like cell spheroids. Prototypical electron dense secretory vesicles (asterisks) are observed dispersed in the cytoplasm. Well-preserved mitochondria (mit), endoplasmic reticulum (ER), Golgi membranes (G) and lipid droplets (LD) are also observed. Inset shows a detail of a secretory vesicle with an average diameter of 450nm. N, nucleus. Scale bars are 200nm (inset) and 500nm. Data are presented as the mean ± SEM for the number of experiments (n) indicated in parenthesis. *,P<0.05; **,P<0.01; ***,P<0.001, relative to 2D iβ cells **(C**,**E)**, or between glucose concentrations **(D**,**F)** using one sample t-test (**C**), unpaired t test **(D, E)** or one-way ANOVA **(F)**.

To establish the glucose threshold for stimulation of insulin secretion, iβ-like cell clusters were subjected to either 2,5,11 or 20mM glucose. Although there was some variability, when moved from 2mM to 5mM glucose, cells did not show a statistical significant increase in insulin secretion (Fig. 5F). They increased insulin secretion on average by a factor of 2.3 when moved from 2mM to 11mM or to 20mM glucose (Fig. 5F). These observations indicate that iβ-like cells are stimulated at higher glucose threshold. Remarkably, this concentration dependence of insulin secretion is similar to human islets that have a threshold at 3mM and a maximal response at 15mM ^39^. In light of these results, we asked whether iβ-like cells presented recognizable secretory granules. Therefore, iβ-like cell clusters where fixed and prepared for conventional electron microscopy. The presence of multiple spherical electron-dense prototypical secretory vesicles was evident in most cells (Fig. 5G). These vesicles showed a high degree of morphological heterogeneity, presumably as consequence of their degree of maturation and/or loading. Although they did not have the appearance of typical insulin-containing granules from primary β cells, which are characterized by a clear halo surrounding a dark polygonal dense core ^42^, some of the vesicles exhibited a grey or less electron dense halo and looked similar to the granules described in insulin-positive cells generated from stem cells in original differentiation protocols ^10, 43^.

### In vivo assessment of human fibroblast-derived iβ-like cells

Lastly, we studied the maintenance of cellular reprogramming in vivo. To this end, we transplanted 300 iβ-like cell spheroids (1000 cells/spheroid) into the anterior chamber of the eye (ACE) of immune-deficient NOD *scid* gamma (NSG) mice (Fig. 6A,B). The ACE allows fast engraftment ^44^ and *in vivo* imaging^45^. To assess *in vivo* graft revascularization we used two-photon microscopy after injection of RITC-dextran and confirmed the presence of functional vessels in the grafts 10 days after transplantation (Fig. 6C and video S1). We also established that the transplanted cells were viable by visualizing fluorescence of the long-term tracer CFDA (Fig. 6C). We then investigated whether reprogramming was retained in vivo. We first studied human *INS* gene expression in eye grafts harvested ten days after transplantation. As shown in Figure 6D, human *INS* transcripts were readily detectable and levels (expressed relative to *TBP*) were similar to those exhibited by iβ-like cells prior to transplantation (Fig.2A, Fig. 5C). Corroborating this result, there were abundant HLA+ (human cell marker) cells that stained for C-peptide and for the transcription factors PDX1, NKX2-2 and MAFA, which are produced from both viral transgenes and cellular genes, in the eye grafts (Fig. 6E and Fig. S5). Of the total number of HLA+ cells counted, 43.5±2.7% were also positive for C-peptide (Fig. 6F). In addition, thirty minutes after an intra-peritoneal glucose injection, human insulin was readily detectable in the aqueous humor of the eyes carrying iβ-like cell grafts (17 of 17, ranging from 11 to 158 mU/L), whereas no insulin was detected in eyes transplanted with non-engineered fibroblast clusters or in mice that received no transplants (Fig.6G). Average insulin levels achieved with 300 iβ-like cells were around 10-fold lower than those observed with 150-200 human islets (53±11 mU/L as compared to 688±99mU/L) (Fig. 6G). Together, these data confirm that cellular reprogramming persisted after transplantation.

**Figure 6.**
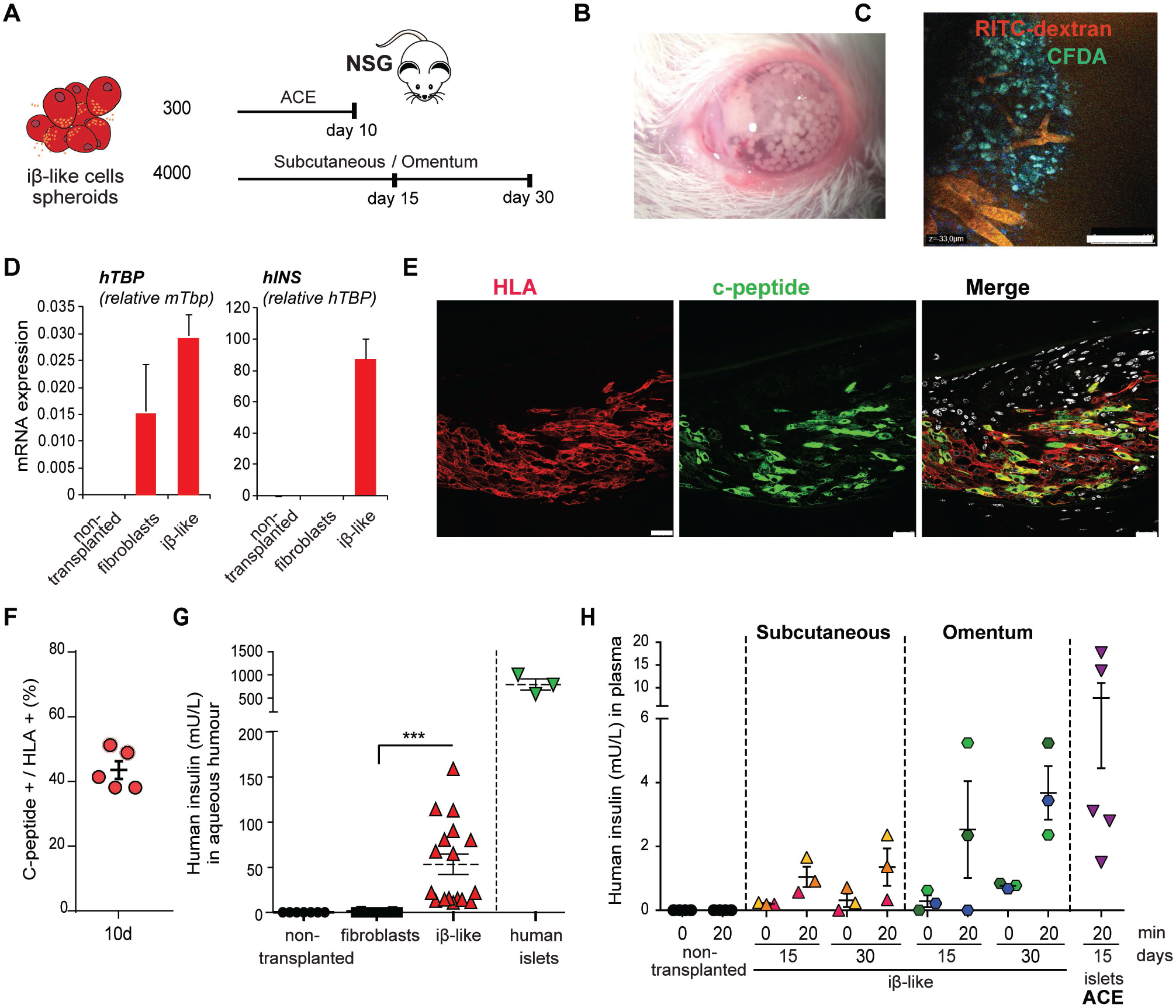
In vivo characterization of iβ-like cell spheroids. **(A)** Schematic representation of the transplantation of iβ-like cell spheroids into the anterior chamber of the eye (ACE) of immunodeficient NSG mice. **(B)** Image of an eye recently transplanted with 300 iβ-like cell spheroids (1000 cells/spheroid). **(C)** Vascularization of iβ-like cell aggregate grafts ten days following transplantation into the ACE. Representative in vivo image of functional vessels (RITC-dextran, red) and viable iβ-like cells (CFDA, green). Scale bar is 100 µm. **(D)** qRT-PCR of human *TBP* and human INS transcripts in eyes of non-transplanted mice (n=3) and mice transplanted with fibroblast spheroids (n=3) or iβ-like cell spheroids (n=5) collected ten days post-transplantation. Human *TBP* expression is calculated relative to mouse*Tbp* and human *INS* relative to human *TBP*. **(E)** Representative immunofluorescence images showing HLA staining in red and C-peptide staining in green in iβ-like cell spheroid grafts ten days post-transplantation in NSG mice. Scale bar is 25 μm. **(F)** Percentage of C-peptide+/HLA+ in eye grafts ten days after transplantation (n=5). **(G)** ELISA determination of human insulin in the aqueous humor after a glucose bolus in un-transplanted mice (n=7), in control mice transplanted with fibroblast spheroids (n=16) and in mice transplanted with iβ-like cell spheroids (n= 17) in the ACE at day 10 post-transplantation; and in mice transplanted with 150-200 human islets (n=3) in the ACE at day 12-15 post-transplantation. **(H)** ELISA determination of human insulin in plasma, before and after a glucose challenge, at days 15 and 30 after transplantation of iβ-like cell spheroids subcutaneously (n=3) or in the omentum (n=3) of NSG mice. Each individual mouse is represented by a different color. Plasma human insulin levels in non-transplanted animals (n=3), before and after glucose administration, and in animals transplanted with human islets in the ACE (n=5), after a glucose bolus, are provided for comparison purposes. Data are presented as mean ± SEM. ***, P< 0.001 using unpaired t-test.

Finally, to evaluate iβ-like cell function in vivo, approximately 4000 spheroids were loaded onto a collagen disk and deposited subcutaneously into the abdominal cavity of adult NSG mice. We detected low levels of circulating human insulin after a glucose challenge in 3 (of 3) and 2 (of 3) mice at days 15 and 30 post-transplantation, respectively (Fig. 6H). We repeated this experiment using the well-vascularized omentum as implant site and observed overall higher human insulin levels and significant glucose-stimulated insulin secretion at day 30 post-transplantation in 3 (of 3) mice (Fig. 6H). Together, these results demonstrate that iβ-like cells exhibit glucose-induced insulin secretion in vivo. Remarkably, glucose-induced plasma human insulin levels observed in mice transplanted with 4000 iβ-like cell spheroids were in the same order of magnitude than mice transplanted with a total of 300-500 human islets into the ACE (Fig. 6H). Thus, correlating with determinations in the aqueous humor, these results reveal that iβ-like cell clusters secrete *in vivo* approximately one tenth of the amount of insulin released by human islets.

## DISCUSSION

In this study, we have developed a 10-day long protocol to produce insulin-producing cells from human fibroblasts through sequential introduction of five lineage-determining transcription factors. Enforced expression of PNM+Pax4+Nkx2-2 results in concomitant activation of the β-cell and down-regulation of the fibroblastic transcriptional programs. Significantly, reprogrammed cells exhibit a functional β-cell like phenotype as judged by their expression of critical β-cell function genes and their ability to mobilize calcium and secrete insulin upon glucose stimulation. These findings warrant further investigations on TF-based reprogramming strategies to generate β-like cells from human fibroblasts.

When compared to other published TF-based reprogramming examples, iβ-like cells exhibited *INS* transcript levels similar to those of reprogrammed human pancreatic duct cells ^20^ and remarkably higher than those reached in human hepatocytes ^46^. Still, expression levels of insulin and of other β-cell marker genes were lower than in human islets. Likewise, glucose-induced insulin secretion was modest, especially in terms of the amount of hormone secreted. We expect further progress by redefining culture conditions as it has been done in stem cell derivation protocols for more than a decade ^7, 10, 43, 47-52^. For example here we report that, by merely moving from 2D to a 3D culture system, iβ-like cells acquired the ability to secrete insulin in response to glucose. Another simple strategy to improve the functionality of iβ-like cells would be the addition of soluble factors during or after introduction of the conversion TFs ^38, 53, 54^. Some of these factors might be selected from the growing body of information on molecules that enhance maturation of β cells generated from stem cells ^55^. Interestingly, it has been shown that soluble molecules can even substitute the function of exogenous factors in some reprogramming protocols ^56-58^.

The use of skin fibroblasts as cell source in place of other less accessible cell lineages can facilitate clinical translation of this approach. Furthermore, the prospect of auto-transplantation and the avoidance of tumor-related concerns linked to pluripotent cell states are two additional assets of this type of direct reprogramming approaches. Here we show that iβ-like cell grafts are efficiently re-vascularized, viable and exhibit glucose-induced insulin secretion in vivo. Circulating human insulin levels in mice transplanted with iβ-like cells were lower but not far-off from those reported in publications using stem cell-derived insulin-producing cells (Table S2). Together, these results provide the foundation for further work aimed at evaluating the long-term function of transplanted iβ-like cell spheroids under pathophysiological contexts. It will be interesting to assess whether the in vivo environment aids in the maturation process of iβ-like cells as it has been reported for stem cell-derived β cell progenitors ^49, 52^.

In summary, our results demonstrate that it is possible to generate insulin-producing cells from human fibroblasts using lineage specific transcription factors. This work evidences the feasibility of direct lineage reprogramming to generate β-like cells from developmentally distant cell lineages. The relative technical simplicity and speed of this procedure together with the opportunity to generate patient-specific β cells for disease modeling and/or autologous transplantation warrants a more comprehensive evaluation of this approach in the future.

## ACKNOWLEDGEMENTS

We are indebted to Lidia Sanchez, Yaiza Esteban (IDIBAPS, CIBERDEM) and Nawelle Dogheche (IDIBAPS) for their help in specific experiments, to Anna Soler (Hospital Clinic) for karyotyping the reprogrammed cells, and to Chris Newgard and Hans Hohmeier (Duke University) for critical reading of the manuscript. We thank the Advanced Optical Microscopy and the Electronic Microscopy Units of the Technological Centers of the University of Barcelona (CCiTUB for their help in sample processing and analyses. The monoclonal antibodies against C-peptide and Nkx2-2 were obtained from the Developmental Studies Hybridoma Bank developed under the auspices of the NICHD and maintained by The University of Iowa. This work has been supported by grant PI19/00896 (to RGa, RGo) integrated in the Plan Estatal de I+D+I and cofinanced by ISCIII-Subdirección General de Evaluación and Fondo Europeo de Desarrollo Regional (FEDER-”A way to build Europe”), grants 120230 (to RGa) and 121430/31/32 (to NM) from La Fundació La Marató de TV3, grant EFSF/JDRF/Lilly Type 1 Programme 2017 (to RGa) from the European Foundation for the Study of Diabetes, grants 2017 SGR 1166 (to JV) and 2017 SGR 1306 (to NM) from the Generalitat de Catalunya, by the Fundación DiabetesCERO and by the Cátedra Astra Zeneca. The research leading to these results has received funding from the European Community’s Seventh Framework Programme (FP7/2009-2013) under the grant agreement nº229673 (SC) and from the European Research Council under the European Union’s Horizon 2020 research and innovation programme-StG-2014-640525_REGMAMKID (NM). CIBERDEM (Centro de Investigación Biomédica en Red de Diabetes y Enfermedades Metabólicas Asociadas) is an initiative of the Instituto de Salud Carlos III.

## AUTHOR CONTRIBUTIONS

Conceived and designed the experiments: MF, RGa. Performed the experiments: MF, AG, EP, NT, HF, RFR, SC. Provided human islets: CB, AW. Provided materials: LC, JR, NM, AN. Analyzed and discussed the data: MF, CE, JV, RGo, RGa. Wrote the manuscript: MF, RGa.

## Supplemental information

**Supplementary S1.**
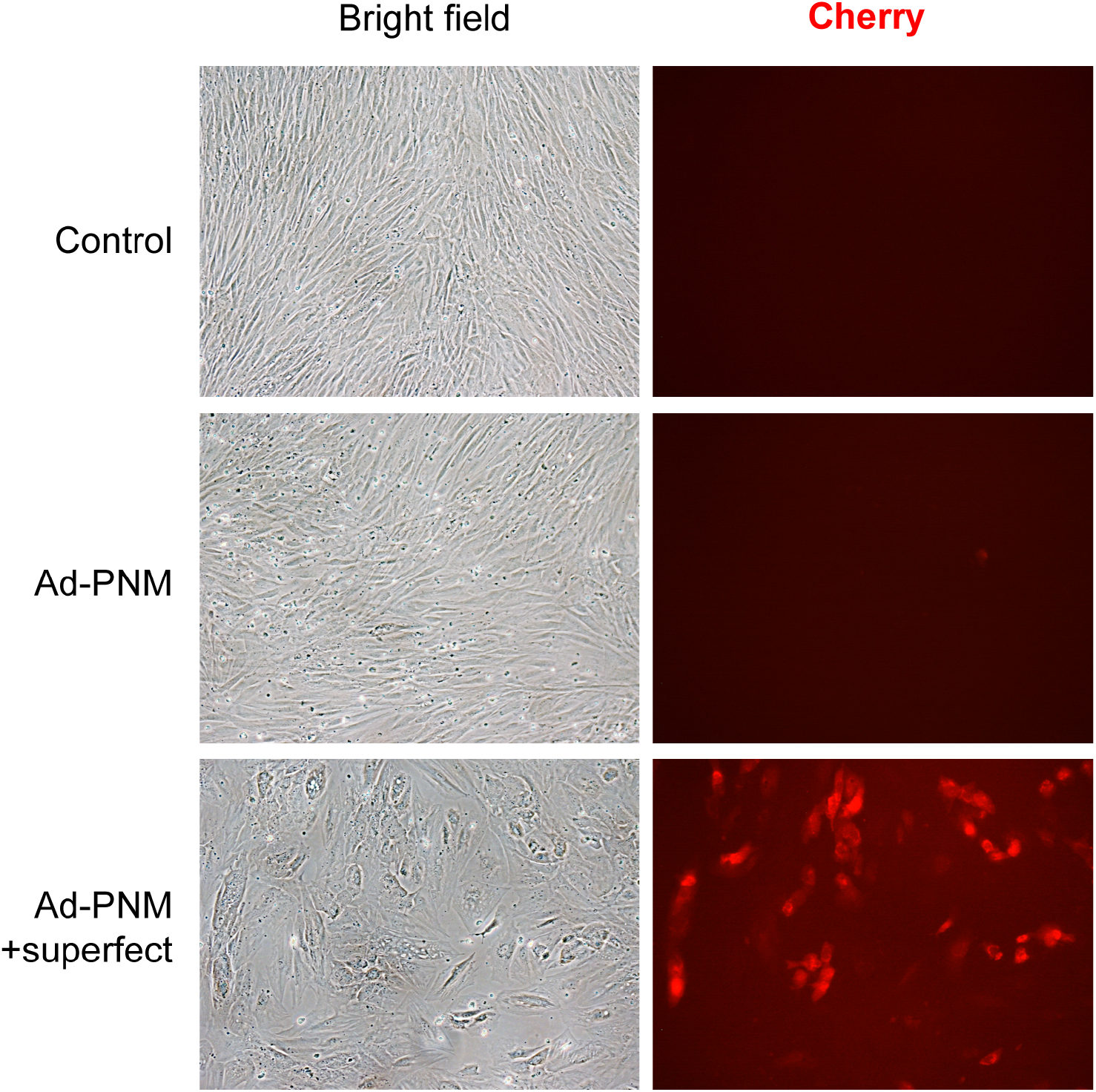
Human primary foreskin fibroblasts HFF1 were treated for 14h with a polycistronic recombinant adenovirus (Ad-PNM) encoding the PNM factors (Pdx1, Neurog3 and Mafa) and a Cherry reporter protein, in the presence or absence of the Superfect transfection reagent. Bright field and Cherry immunofluorescence images from control (parental) fibroblasts and HFF1 fibroblasts four days after infection.

**Supplementary S2.**
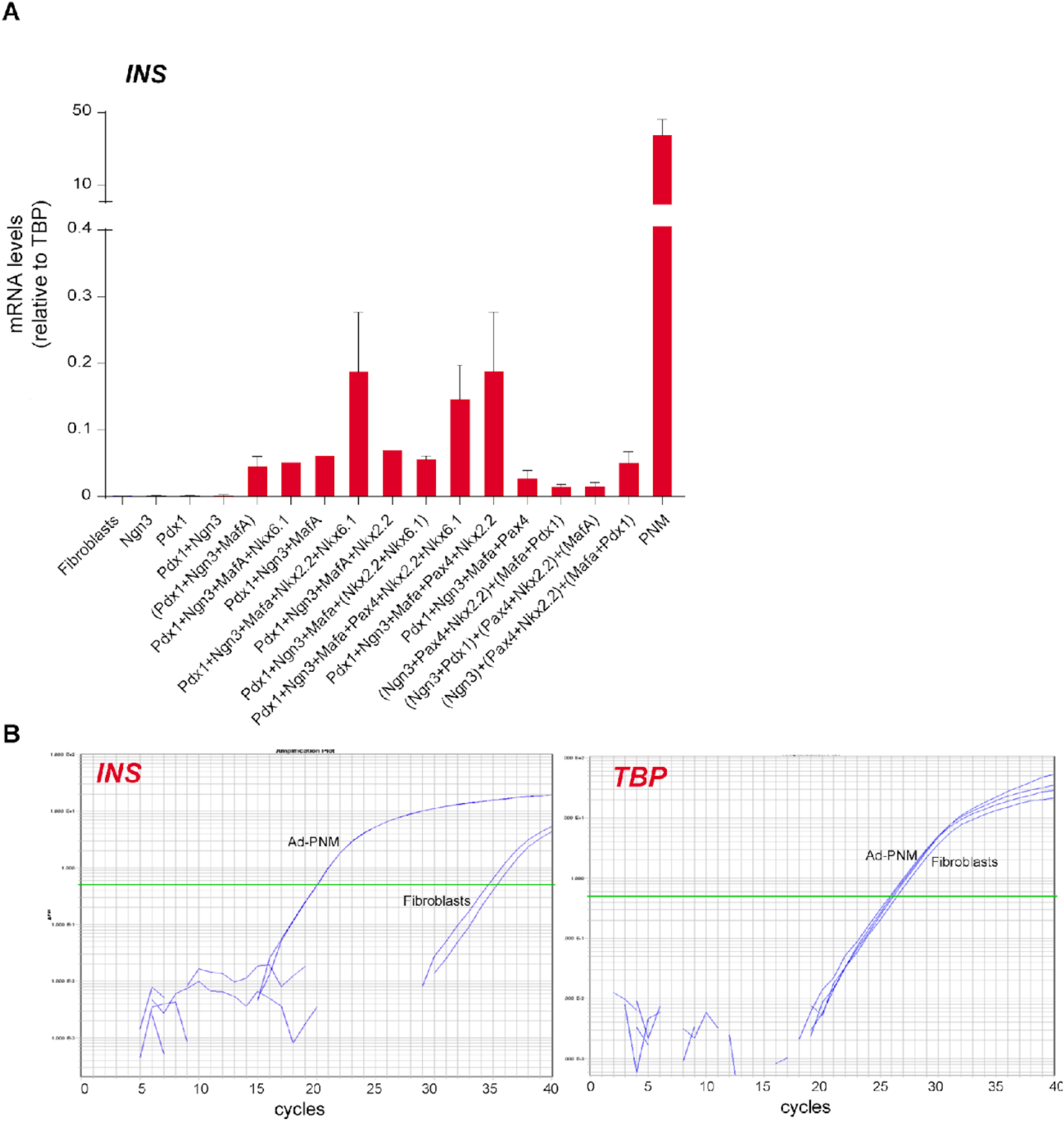
**(A)** qRT-PCR for human *INS* in human foreskin fibroblasts HFF1 ten days after infection with the recombinant adenoviruses encoding the indicated transcription factors. All adenoviruses except Ad-PNM encoded a single transcription factor. Adenoviruses that were added simultaneously are shown in parenthesis. Order of addition is left to right. Expression levels were calculated relative to *TBP*. Data are mean ± SEM for n=2-9. **(B)** Real time amplification plots for *INS* and *TBP* in parental fibroblasts and in fibroblasts seven days after infection with Ad-PNM.

**Supplementary S3:**
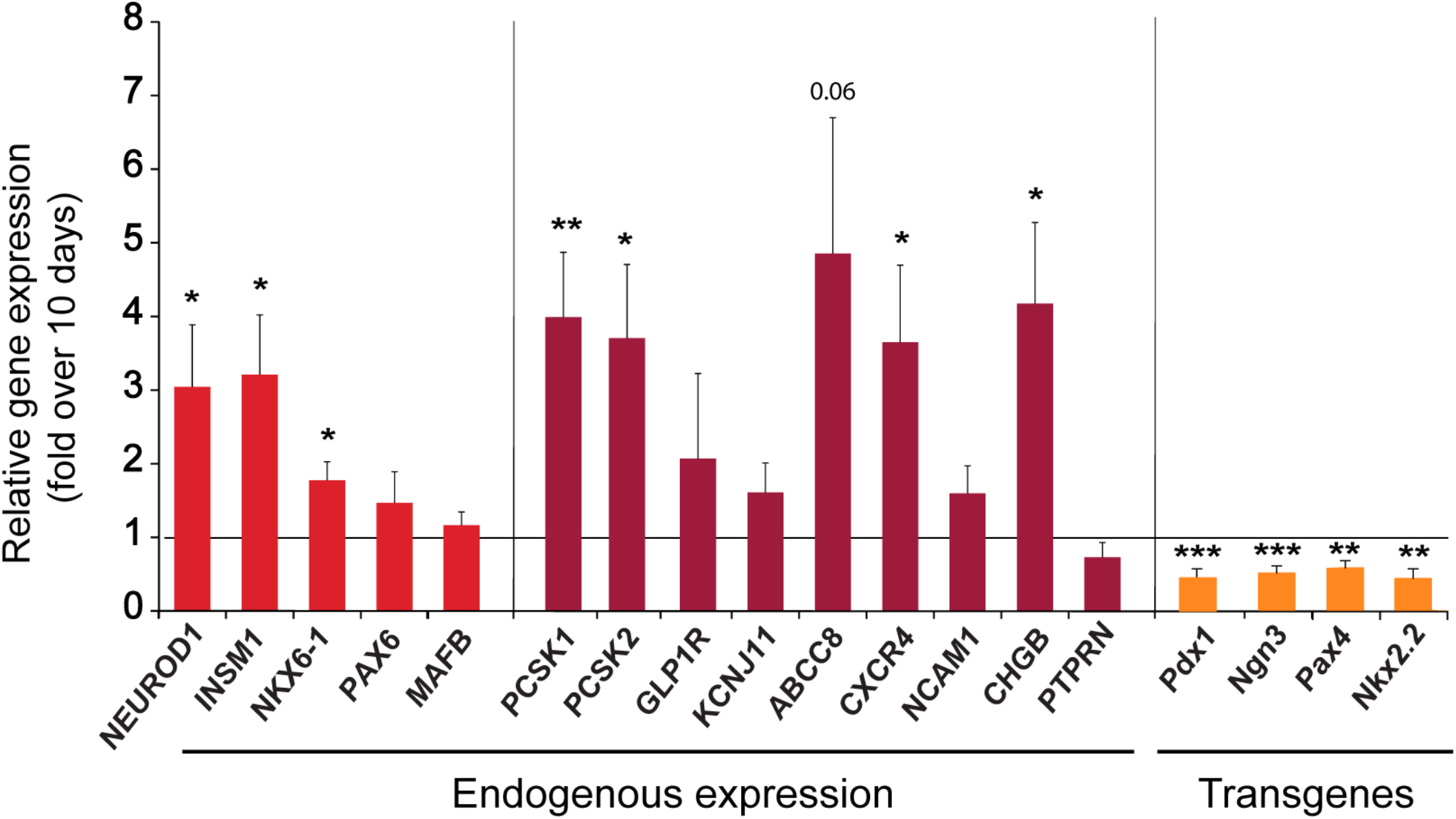
qRT-PCR of the indicated genes at day 21 after initiation of reprogramming. Transcript levels are expressed relative to levels in cells at day 10 of the reprogramming protocol (given the value of 1). Cells were maintained in 2D culture throughout the duration of the experiment. Data are mean ± SEM for n=9-13, from 7 reprogramming experiments. *P< 0.05, **P< 0.01, ***, P< 0.001, vs. iβ-cells at day10 using

**Supplementary S4:**
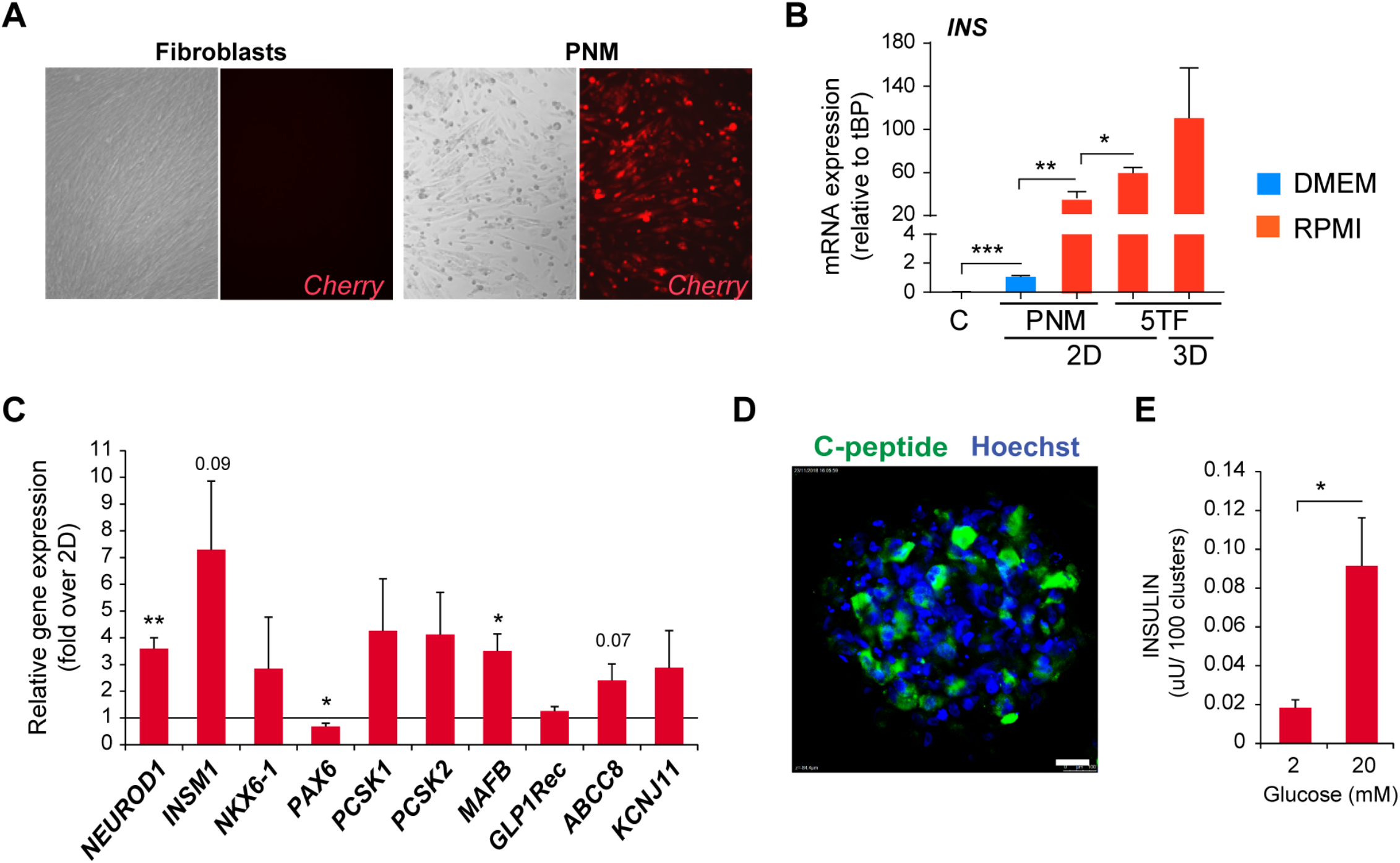
Human primary foreskin fibroblasts HFF2 were subjected to transcription factor-based reprogramming towards insulin-producing cells. **(A)** Bright field and Cherry immunofluorescence images from control (parental) fibroblasts and from fibroblasts three days after infection with Ad-PNM. Scale bar is 100 uM. **(B)** qRT-PCR for *INS* in HFF2 ten days after infection with Ad-PNM alone and cultured under 2D conditions in DMEM (blue) or RPMI-1640 media (red) (n=6), or subjected to the 5TF reprogramming protocol and cultured in RPMI-1640 media under 2D or 3D (n=4) conditions as indicated. Expression levels were calculated relative to *TBP*. Note that changes in *INS* gene expression in the different reprogramming conditions follow a similar trend as observed with HFF1 fibroblasts. **(C)** qRT-PCR of the indicated genes in HFF2-derived iβ-like cell spheroids (n=6) Transcript levels are expressed relative to HFF2-derived iβ-like cells maintained in 2D culture throughout the 10-day protocol (given the value of 1). **(D)** Representative immunofluorescence image showing C-peptide staining in green and nuclei in blue (marked with Hoechst). Scale bar is 100 µm. **(E)** In vitro glucose-induced insulin secretion by iβ-like cell spheroids. Data are calculated as mean ± SEM for 3 independent experiments, 1-3 replicates per experiment. *, P< 0.05; **, P< 0.01; ***, P< 0.001 relative to 2D in C using one-sample test, or between indicated conditions in B and E using unpaired t-test.

**Supplementary S5:**
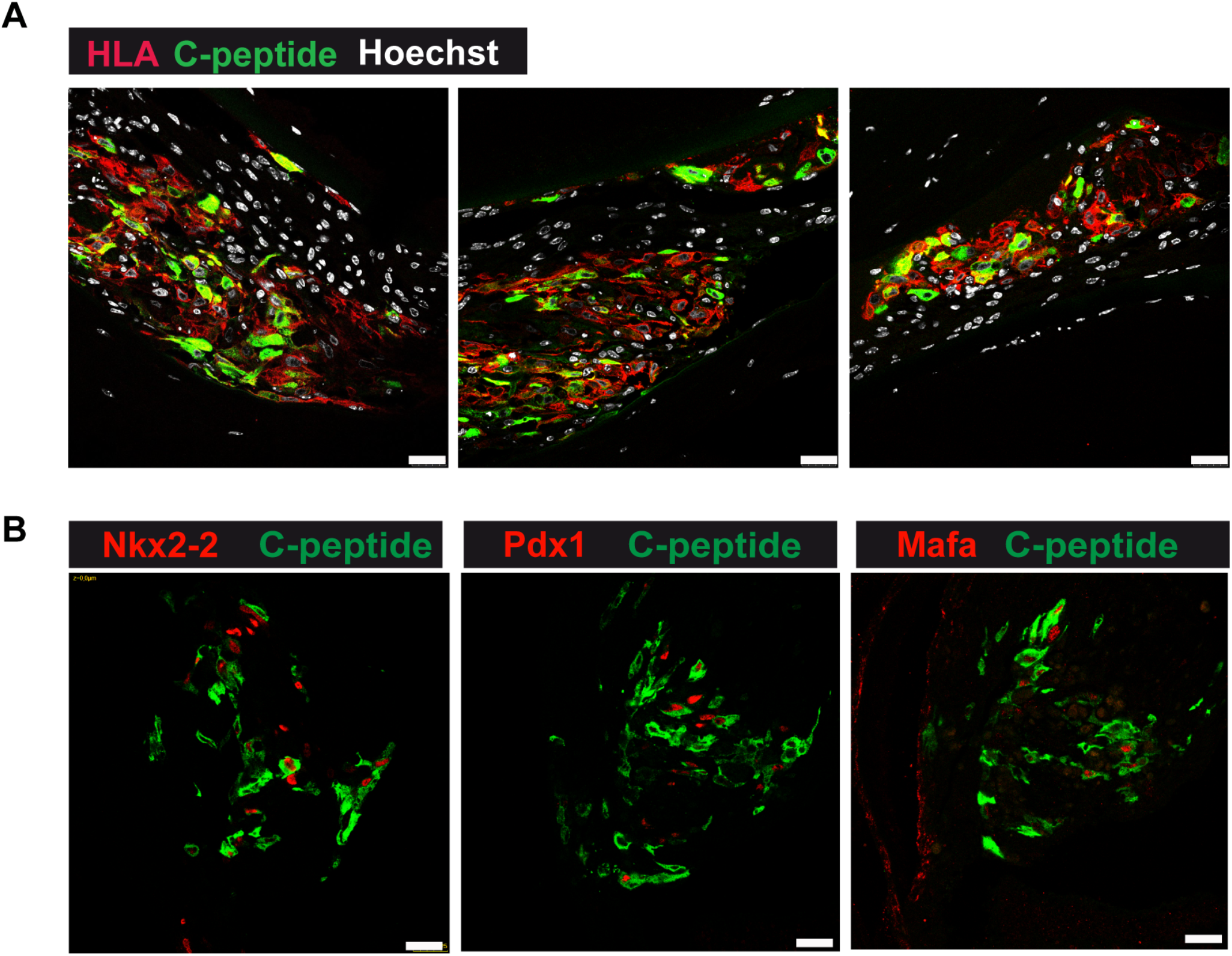
Immunostaining of iβ-cell spheroids ten days after their transplantation into the anterior chamber of the eye of NSG mice. Representative immunofluorescence images showing **(A)** C-peptide staining (green), HLA (red) and cell nuclei marked with Hoechst (white) and **(B)** C-peptide (green) with the indicated transcription factors (red). Scale bars are 25 µm.

**Table S1.** Comparison of gene expression levels between iβ-cells maintained in 2D culture, iβ-cell spheroids (3D) and human islets for the indicated genes. Fold expression values are calculated using qPCR data presented in Figures 1D&E, 2A, 3A-C and 5C.

| **Gene** | **Fold expression**<br>**Islets/ iβ-cells 2D** | **Fold expression**<br>**Islets/ iβ-cell 3D** |
| --- | --- | --- |
| ***ABCC8*** | 2335 | 450 |
| ***CHGB*** | 23036 | 9251 |
| ***CX36*** | 3589 | 399 |
| ***CXCR4*** | 6 | 2 |
| ***GIPR*** | 10 | 6 |
| ***GLP1R*** | 163 | 100 |
| ***INS*** | 358 | 205 |
| ***INSM1*** | 134 | 41 |
| ***KCNJ11*** | 51 | 12 |
| ***MAFB*** | 8 | 4 |
| ***NCAM1*** | 2 | 2 |
| ***NEUROD1*** | 200 | 63 |
| ***NEUROG3*** | 2 | 3 |
| ***NKX2-2*** | 196 | 265 |
| ***NKX6-1*** | 117 | 37 |
| ***PAX4*** | 14 | 9 |
| ***PAX6*** | 425 | 327 |
| ***PCSK1*** | 16 | 5 |
| ***PCSK2*** | 81 | 48 |
| ***PDX1*** | 115 | 68 |
| ***PTPRN (IA2)*** | 30 | 25 |

**Table S2.**
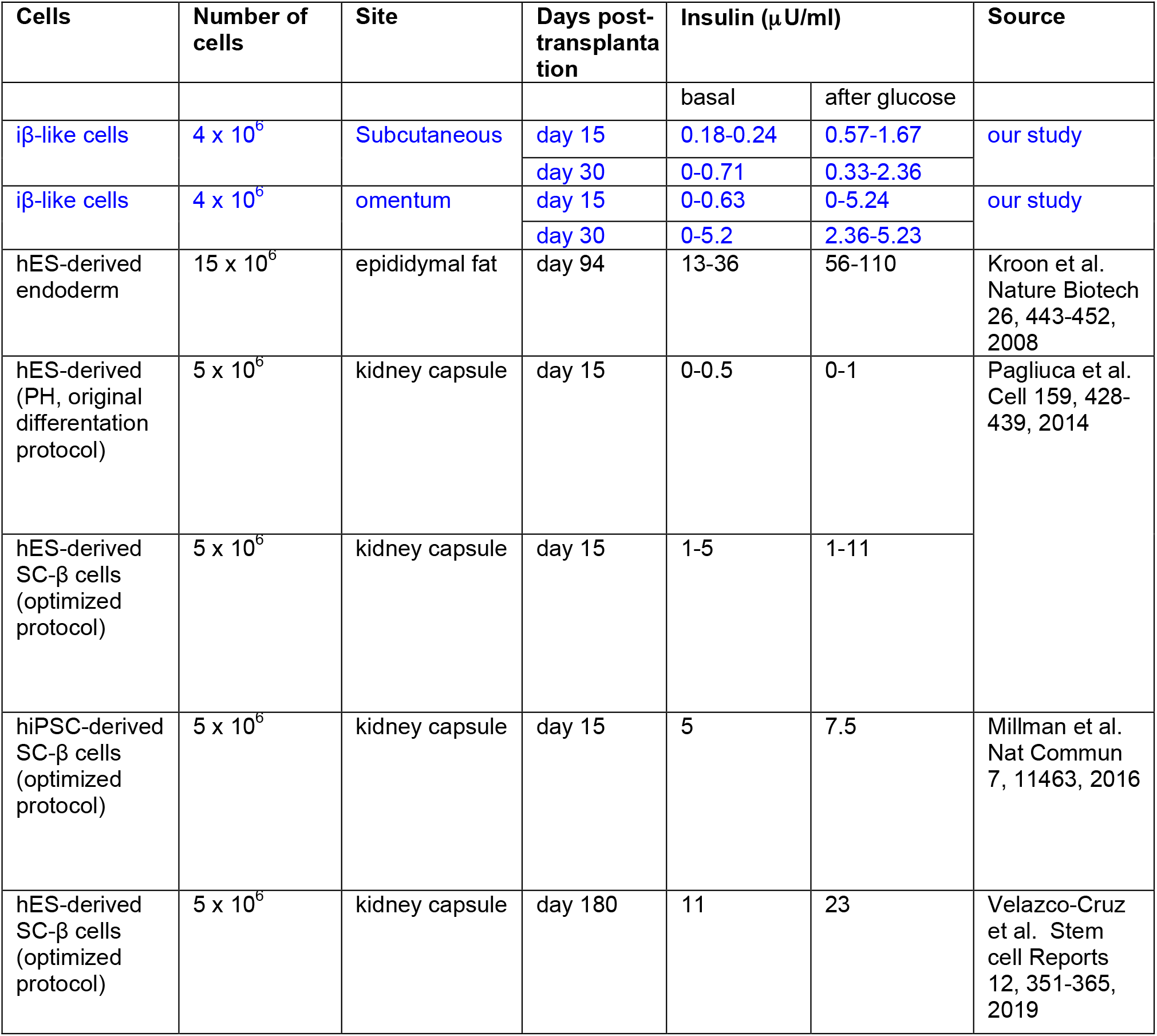
Comparison of circulating human insulin levels in transplanted mice between our study and several publications using stem cell-derived β-like cells.

| Cells | Number of cells | Site | Days post-transplantation | Insulin (μU/ml) | | Source |
| --- | --- | --- | --- | --- | --- | --- |
|  |  |  |  | basal | after glucose |  |
| iβ-like cells | 4 x 10<sup>6</sup> | Subcutaneous | day 15 | 0.18-0.24 | 0.57-1.67 | our study |
|  |  |  | day 30 | 0-0.71 | 0.33-2.36 |  |
| iβ-like cells | 4 x 10<sup>6</sup> | omentum | day 15 | 0-0.63 | 0-5.24 | our study |
|  |  |  | day 30 | 0-5.2 | 2.36-5.23 |  |
| hES-derived endoderm | 15 x 10<sup>6</sup> | epididymal fat | day 94 | 13-36 | 56-110 | Kroon et al. Nature Biotech 26, 443-452, 2008 |
| hES-derived (PH, original differentiation protocol) | 5 x 10<sup>6</sup> | kidney capsule | day 15 | 0-0.5 | 0-1 | Pagliuca et al. Cell 159, 428-439, 2014 |
| hES-derived SC-β cells (optimized protocol) | 5 x 10<sup>6</sup> | kidney capsule | day 15 | 1-5 | 1-11 | |
| hiPSC-derived SC-β cells (optimized protocol) | 5 x 10<sup>6</sup> | kidney capsule | day 15 | 5 | 7.5 | Millman et al. Nat Commun 7, 11463, 2016 |
| hES-derived SC-β cells (optimized protocol) | 5 x 10<sup>6</sup> | kidney capsule | day 180 | 11 | 23 | Velazco-Cruz et al. Stem cell Reports 12, 351-365, 2019 |

**Table S3.**
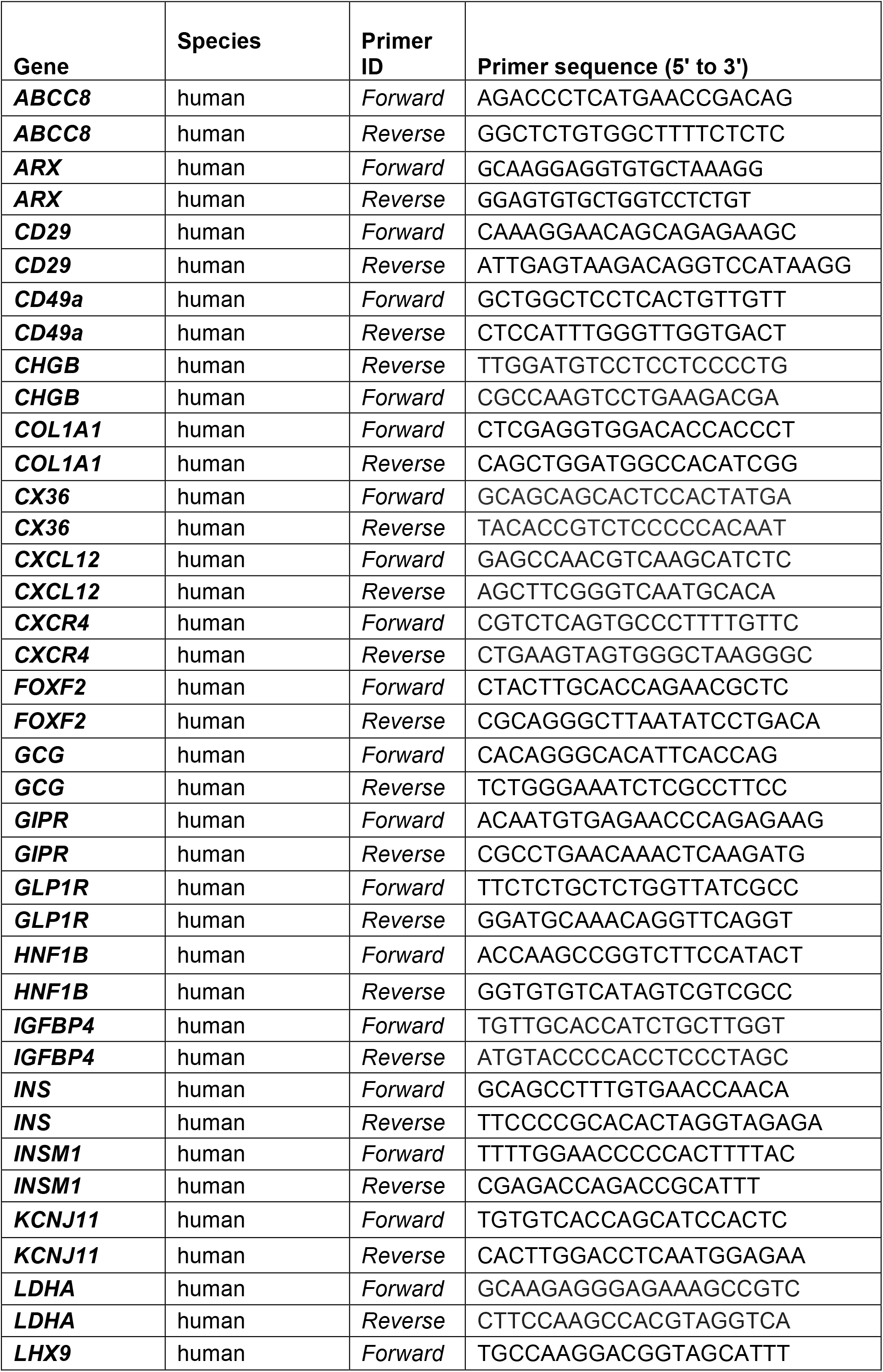
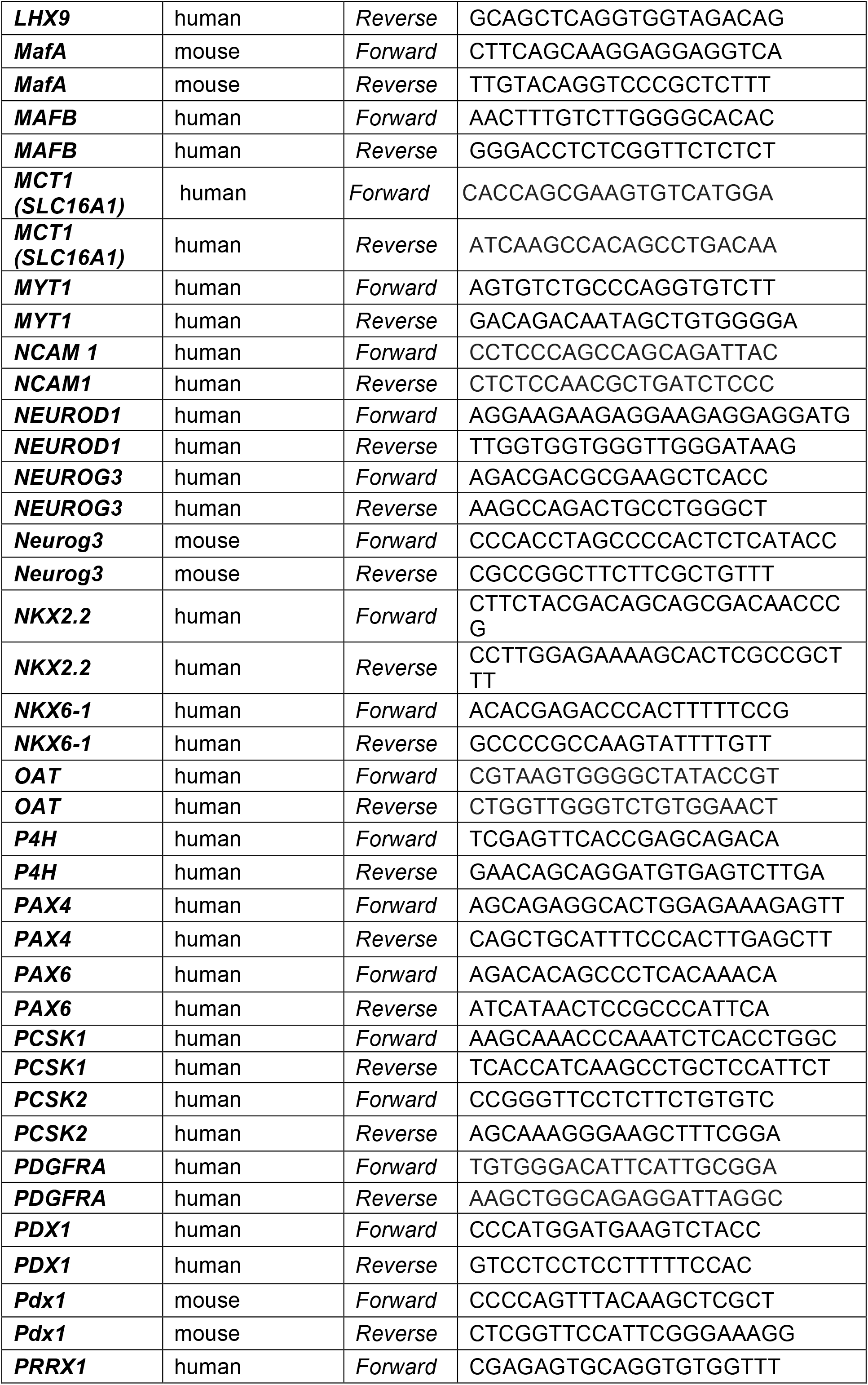
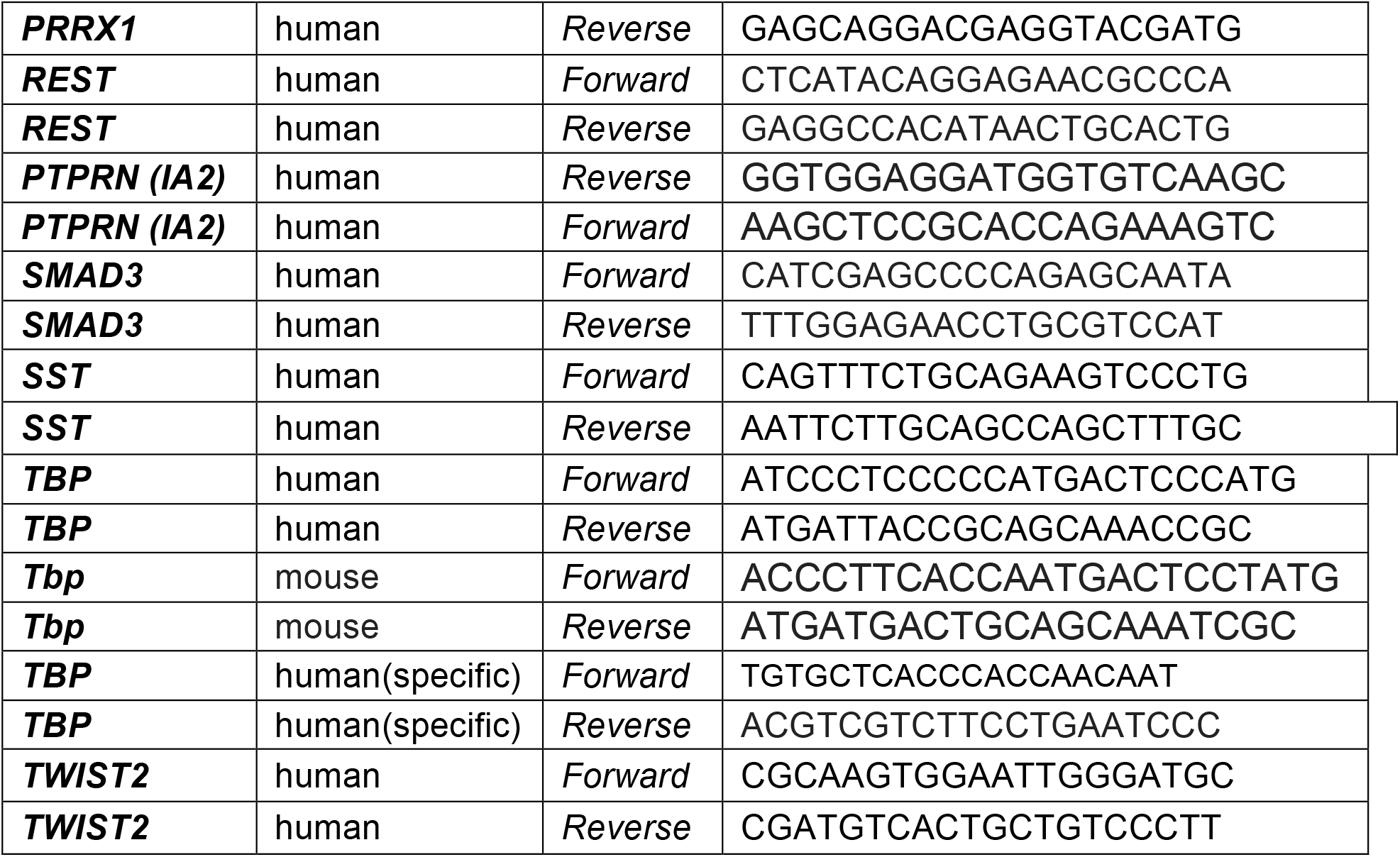
List of oligonucleotides used for real time PCR.

## VIDEO CAPTIONS

**Video S1**

Video shows changes in fluorescence of an isolated iβ-like cell in response to high glucose (20mM) and high potassium (KCl 30mM). Cell had been pre-loaded with the calcium indicator Fluo-2. Video was recorded using a Leica TCS SPE confocal microscope with an incubation chamber set at 37°C, and a 40X oil immersion objective.

**Video S2**

Video shows fluorescence of parental fibroblasts HFF1 in response to high glucose (20mM) and high potassium (KCl 30mM). Cells had been pre-loaded with the calcium indicator Fluo-2. Note that fibroblasts exhibit no changes in fluorescence in response to the tested stimuli. Video was recorded using a Leica TCS SPE confocal microscope with an incubation chamber set at 37°C, and a 40X oil immersion objective.

